# ANXA2 enhances virus replication through negatively regulating cGAS-STING and RLRs-mediated signal pathway

**DOI:** 10.1101/2021.12.01.470696

**Authors:** Mengdi Xue, Hongyang Liu, Zhaoxia Zhang, Chunying Feng, Kunli Zhang, Xiaohong Liu, Guangqiang Ye, Qiongqiong Zhou, Changyao Li, Changjiang Weng, Li Huang

## Abstract

Host nucleic acid receptors can recognize the viral DNA or RNA upon virus infection, which further triggers multiple signal pathways to promote the translocation of the interferon regulatory factor 3 (IRF3) into nucleus and produce type I interferon (IFN), leading to the host antiviral response. Here, we report a novel negative regulator Annexin A2 (ANXA2) that regulates type I IFN production through multiple mechanisms. Ectopic expression of ANXA2 inhibited the production of type I IFN induced by DNA- and RNA viruses and enhanced virus replication, while knockout of ANXA2 expression enhanced the production of type I IFN and inhibited virus replication. Mechanistically, ANXA2 not only disrupted MDA5 recruiting MAVS, but also inhibited the interaction between MAVS and TRAF3 upon RNA virus infection. In addition, ANXA2 impacted the translocation of STING from endoplasmic reticulum to Golgi apparatus upon DNA virus infection. Interestingly, ANXA2 also inhibited IRF3 phosphorylation and nuclear translocation through competing with TANK-binds kinase 1 (TBK1) and inhibitor-κB kinase ε (IKKε) for binding to IRF3. *Anxa2* deficiency *in vivo* increased the production of type I IFN, which resulted in suppression of encephalomyocarditis virus (EMCV) replication. Our findings reveal that ANXA2, as a negative regulator of type I IFN production, plays an important role in regulating the host antiviral responses.

**Author summary:** Annexin is a family of evolutionarily conserved multi-gene proteins, which are widely distributed in various tissues and cells of plants and animals. These proteins can reversibly bind to phospholipid membranes and to calcium ions (Ca^2+^). To date, several studies have confirmed that some members of the Annexin family regulate the antiviral innate immune response. Until now, regulation of the production of type I IFN by ANXA2 is not reported. In this study, ANXA2 were found to strongly inhibit the production of type I IFN, leading to increased virus replication while knockout of ANXA2 expression inhibited virus replication by increasing the amount of IFN. Compared with wild-type littermates, ANXA2 deficiency mice produced more type I IFN to inhibit virus replication. Our results provide methanistic insights into the novel role of ANXA2 in the antiviral innate immune responses.

## Introduction

Innate immunity, also known as non-specific immunity or natural immunity, is the first line of defense against pathogenic microorganism infection and plays a key role in regulating host antiviral responses to eliminate pathogens. Upon a pathogen infection, the pattern recognition receptors (PRRs) can recognize the conserved molecular structures of the pathogen called pathogen-associated molecular patterns (PAMPs), including viral DNA, viral RNA and surface glycoproteins[1]. The known PRRs include the Toll-like receptors (TLRs), the retinoic acid-inducible gene I (RIG-I)-like receptors (RLRs), the nucleotide-binding domain and leucine-rich repeat-containing receptors (NLRs), the C-type lectin receptors (CLRs), and the PYRIN family members[2].

It is well known that RIG-I and MDA5 are well-conserved RLRs that recognize viral RNAs in the process of RNA virus infection[3]. Although RIG-I and MDA5 play non-redundant roles by detecting different RNA viruses and by recognizing distinct features of viral RNAs, they share the same downstream adaptor molecule, MAVS (also known as IPS-1, VISA or Cardif). Upon RNA virus infection, the carboxy-terminal domain of RIG-I/MDA5 interacts with viral RNA to promote conformational changes of RIG-I/MDA5, which promote them to oligomerize[3] and expose their own caspase activation recruitment domain (CARD). RIG-I/MDA5 CARD interacts with the CARD domain of a mitochondrial antiviral signal protein, MAVS, resulting in recruiting TRAFs to the linker between CARD and TM of MAVS[4], which in turn activates TBK1, IKKε[5], resulting in the phosphorylation and translocation of IRF3 to nucleus[6] to produce type I IFN.

Upon DNA virus infection, the cyclic GMP-AMP (cGAMP) synthase (cGAS) senses the viral DNA and catalyzes ATP-GTP transformation to produce the second messenger 2’3’-cGAMP, which further binds to stimulator of interferon genes (STING). Subsequently, the STING travels from the endoplasmic reticulum (ER) to the ER-Golgi intermediate compartment (ERGIC) and the Golgi apparatus[7–9]. During this process, the carboxy-terminal of STING recruits and activates the TBK1, then the activated TBK1 phosphorylates IRF3[10–13]. The phosphorylated IRF3 forms a homodimer, which is then translocated to the nucleus and binds to the interferon stimulus response element (ISRE) of the target genes to induce type I IFN production[14–16].

Annexin A2 (ANXA2, also known as calpactin I or lipocortin II) is a multifunctional protein that can reversibly bind to phospholipid membranes and calcium ions (Ca^2+^)[17]. ANXA2 belongs to the membrane scaffold and binding protein, and is mainly expressed in the plasma membrane and intracellular vesicles[18, 19]. Although ANXA2 exists in the cytoplasm as a monomer, it generally exists as a heterotetrameric complex composed of S100A10 dimer and two ANXA2 molecules[20, 21]. It has been reported that ANXA2 participates in the replication process of a variety of viruses. For examples, ANXA2 interacts with the matrix (M) protein of viruses (such as measles virus and bovine transient fever virus), which is located on the inner surface of the virus envelope and plays a role in the formation of viral particles, thereby contributing to the assembly and release of the virus[22, 23]. ANXA2 also participates in Classic swine fever virus (CSFV) and Hepatitis C virus (HCV) production process by binding to the NS5A protein[24]. ANXA2 can induce the formation of lipid raft microdomains by recruiting the HCV NS protein and enriching it in lipids, which promotes the formation of the HCV replication complex[25]. In addition, ANXA2 interacts with viral-encoded proteins such as avian influenza virus (AIV) and porcine reproductive and respiratory syndrome virus (PRRSV) to enhance virus replication[26, 27]. However, the role of ANXA2 in regulating type I IFN production remains poorly understood.

In this study, we identified ANXA2 as a negative regulator in the type I IFN production We presented biochemical and genetic evidence that ANXA2 regulated type I IFN production through multiple mechanisms. ANXA2 not only inhibited MDA5 recruiting MAVS and MAVS recruiting TRAF3 upon RNA virus infection, but also inhibited the translocation of STING to Golgi upon HSV-1 infection. In addition, ANXA2 inhibited IRF3 phosphorylation and nuclear translocation by disrupting the interaction among IRF3, TBK1 and IKKε, resulting in inhibition of the type I IFN production and enhancement of viral replication.

## Results

### ANXA2 inhibits type I IFN production

To explore the function of ANXA2 in host antiviral responses, we firstly tested the expression of ANXA2 upon virus infection. The results showed that both RNA virus (vesicular stomatitis virus (VSV), EMCV) and DNA virus (Herpes simplex virus 1 (HSV-1)) induced ANXA2 expression (Fig S1), suggesting that ANXA2 may be involved in host antiviral responses. Further study showed that human ANXA2 inhibited the activation of the IFN-β, IRF7, ISG54, ISRE and NF-κB promoters induced by Sendai virus (SeV) in a dose-dependent manner (Fig 1A and Fig S2A-2D). Similarly, ectopically expressed ANXA2 significantly inhibited the protein level of the IFN-β induced by SeV (Fig 1B). Additionally, the inhibitory effects of ectopic expressed ANXA2 on the mRNA levels of *Ifnβ1* and *Isg56* were observed after transfection with poly(I:C), cGAMP or infection with other RNA viruses, such as EMCV, VSV, and DNA virus (HSV-1) (Fig 1C and 1D).

**Figure 1.**
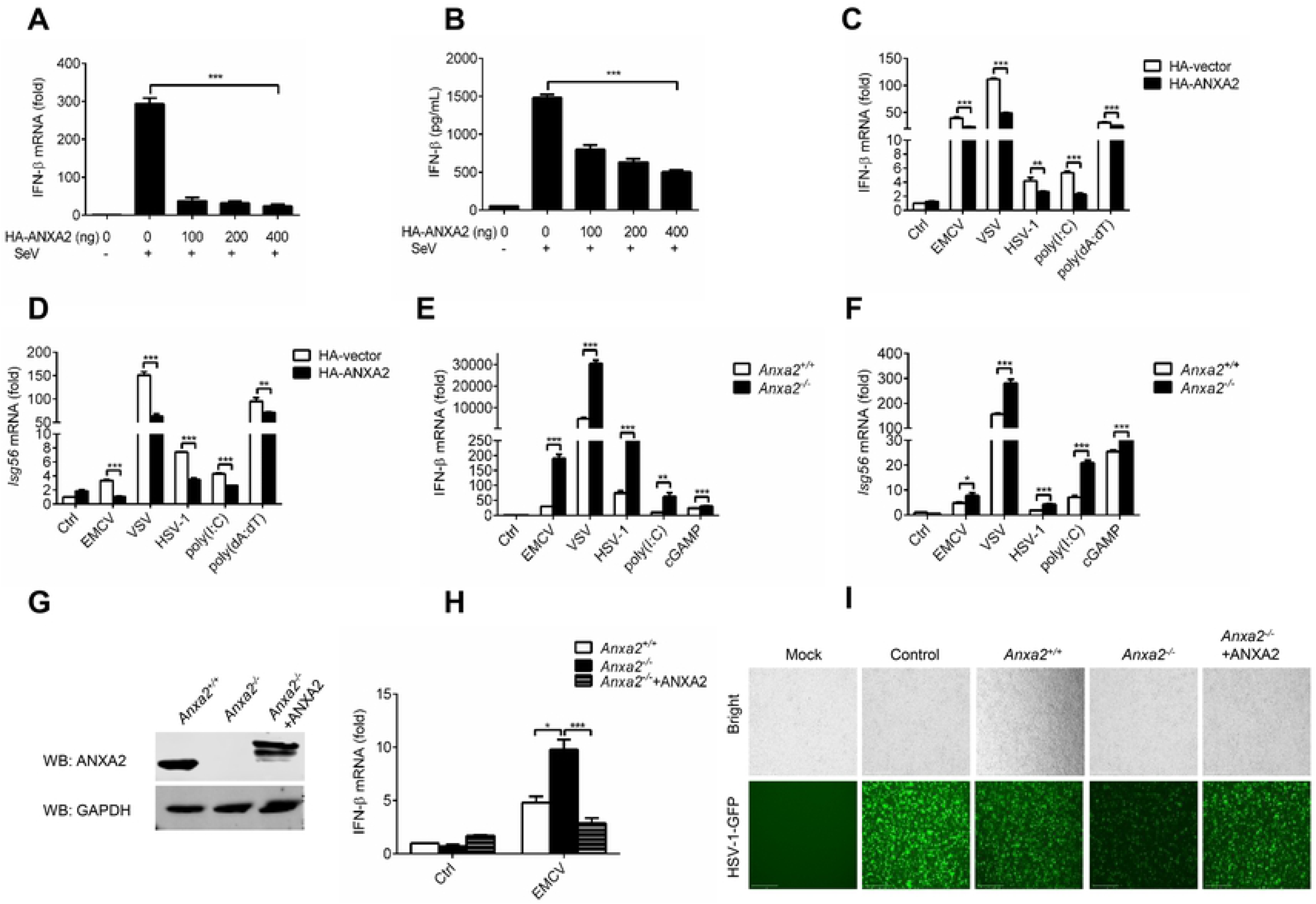
ANXA2 inhibits type I IFN production. (A and B) ELISA and qPCR analysis of the protein levels of IFN-β in the cell culture supernatant and mRNA levels of *Ifnβ1* in HEK293T cells transfected with increasing amount of plasmids expressing HA-ANXA2 and then infected with SeV (1 multiplicity of infection (MOI)) for 12 h. (C and D) qPCR analysis of the mRNA levels of *Ifnβ1* (C) and *Isg56* (D) in the HEK293T cells transfected with HA-vector or HA-ANXA2 and infected with EMCV, VSV or HSV-1 for 12 h or transfected with poly(I:C) (2 μg/mL) or poly(dA:dT) (2 μg/mL) for 24 h. (E and F) qPCR analysis of the mRNA levels of *Ifnβ1* (E) and *Isg56* (F) in HeLa-*Anxa2^+/+^* and HeLa-*Anxa2^−/−^* cells after infection with EMCV, VSV or HSV-1 for 12 h or transfected with poly(I:C) or cGAMP for 24 h. (G) Immunoblot analysis of ANXA2 expression in the HeLa-*Anxa2^+/+^*cells, HeLa-*Anxa2^−/−^* cells and HeLa-*Anxa2^−/−^* cells transfected with a plasmid expressing HA-ANXA2. (H) qPCR analysis of the mRNA levels of *Ifnβ1* in the HeLa-*Anxa2^+/+^* cells, HeLa-*Anxa2^−/−^* cells and HeLa-*Anxa2^−/−^* cells transfected with a plasmid expressing HA-ANXA2 with or without EMCV infection for 12 h. (I and J) HeLa-*Anxa2^+/+^* cells and HeLa-*Anxa2^−/−^* cells transfected with an empty vector or a plasmid expressing ANXA2 were infected with SeV for 12 h. The cell supernatants were placed under ultraviolet (UV) aseptically irradiated for 12 h and then added to HEK293T cells and incubated for 24 h, and the HEK293T cells were infected with VSV-GFP. After 12 h, the replication of VSV-GFP was analyzed using a fluorescence microscope.

To further confirm these results, HEK293T cells with ANXA2 gene deletion (HEK293T-*Anxa2^−/−^*) were generated using CRISPR/Cas9 to examine the function of ANXA2. As shown in Fig 1E and 1F, ANXA2 deficiency markedly increased the mRNA levels of *Ifnβ1* and *Isg56* induced by infection with EMCV, VSV and HSV-1 or transfection with poly(I:C) or cGAMP. Of note, HEK293T-*Anxa2^−/−^*cells transfected with a plasmid expressing ANXA2 significantly reduced the mRNA expression levels of *Ifnβ1* which were enhanced by ANXA2 deficiency (Fig 1G and 1H). An IFN sensitivity result showed that the replication levels of the HSV-1-GFP and VSV-GFP correlated with the expression level of ANXA2 (Fig 1I, Fig S2E and 2F). Taken together, our findings demonstrated that ANXA2 inhibits type I IFN production.

### ANXA2 deficiency enhances cellular antiviral responses *in vitro*

To study the function of ANXA2 in type I IFN production *in vivo*, we generated *Anxa2* knock out (*Anxa2^−/−^*) mice using a homologous recombination technique and validated them by sequencing and Western blot analysis (Fig S3A-C). Primary peritoneal macrophages isolated from *Anxa2^−/−^* mice and their wildtype (WT) littermates were infected with EMCV, VSV or HSV-1 for 0, 4, 8, and 12 h. Compared to macrophages from WT mice, macrophages from *Anxa2^−/−^* mice infected with these viruses showed higher mRNA expression of *Ifnβ1* (Fig 2A-C), *Isg56* (Fig 2D-F) and *Mx1* (Fig 2G-I) at different time points.

**Figure 2.**
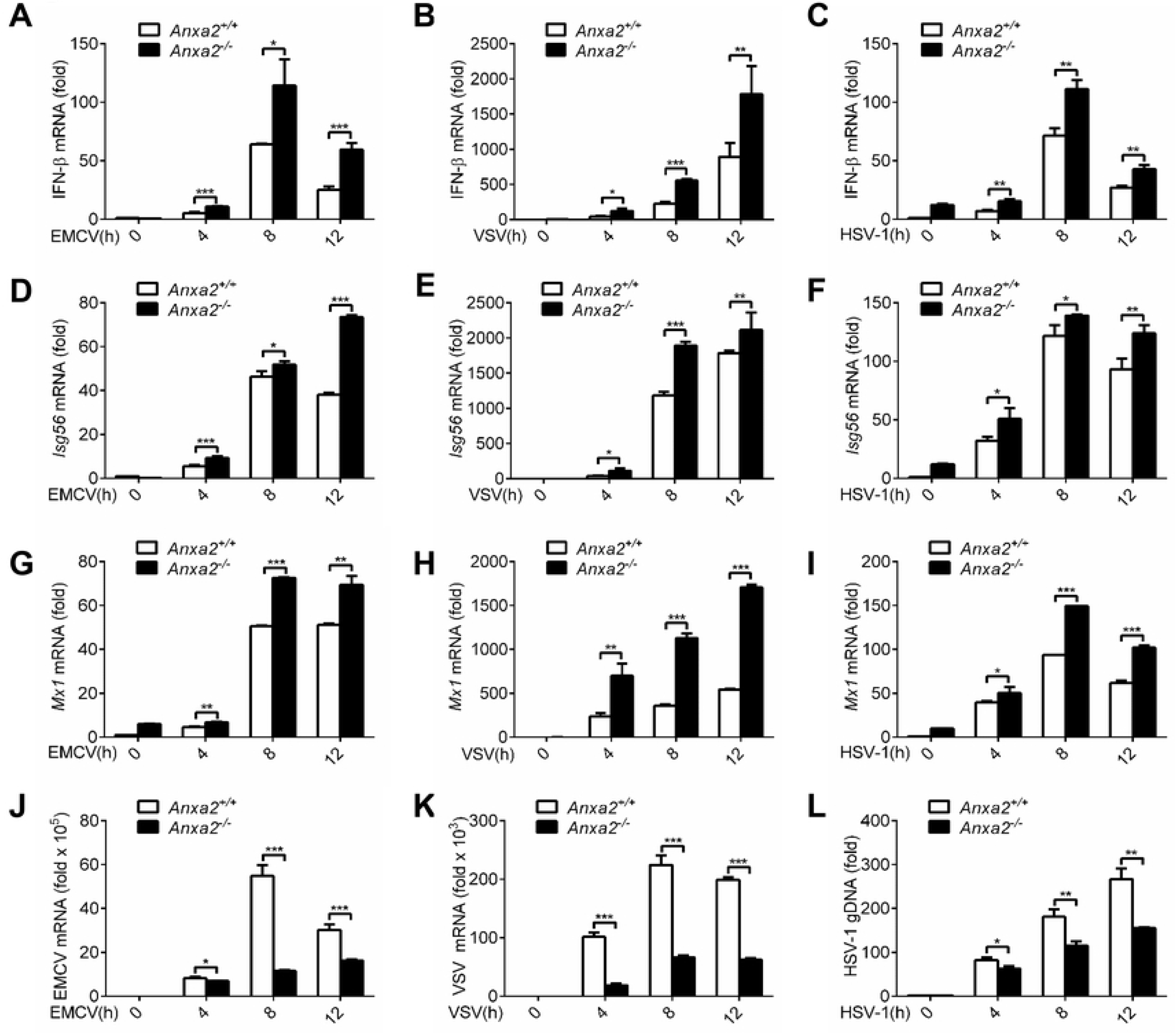
ANXA2 deficiency enhances cellular antiviral responses. (A-C) qPCR analysis of the mRNA levels of *Ifnβ1* in *Anxa2^+/+^* and *Anxa2^−/−^* peritoneal macrophages infected with EMCV (2 MOI) (A), VSV (0.2 MOI) (B), and HSV-1 (10 MOI) (C) for 0, 4, 8 or 12 h. (D-F) qPCR analysis of the mRNA levels of *Isg56* in peritoneal macrophages isolated from the *Anxa2^+/+^* and *Anxa2^−/−^* mice infected with EMCV (D), VSV (E), and HSV-1(F) for 0, 4, 8 or 12 h. (G-I) qPCR analysis of the mRNA levels of *Mx1* in the peritoneal macrophages isolated from *Anxa2^+/+^* and *Anxa2^−/−^* mice infected with EMCV (G), VSV (H), and HSV-1 (I) for 0, 4, 8 or 12 h. (J-L) qPCR analysis of the genomic copy numbers of EMCV (J), VSV (K) or HSV-1 (L) in the peritoneal macrophages isolated from *Anxa2^+/+^* and *Anxa2^−/−^* mice infected with EMCV, VSV or HSV-1 for 0, 4, 8 or 12 h. \**P* < 0.05, \*\**P* < 0.01 and \*\*\**P* < 0.001 (two-tailed Student’s t-test (A-L). Data are representative of three independent experiments with three biological replicates (mean ± s.d. in A-L).

Furthermore, the VSV and EMCV genomic RNA copy number and the HSV-1 genomic DNA copy number in peritoneal macrophages from *Anxa2^−/−^* mice were significantly lower than that of the primary peritoneal macrophages from WT mice at 0, 4, 8, and 12 hpi (Fig 2J-L). Consistent with these results, the mRNA levels of *Ifnβ1*, *Isg56* and *Mx1* in the HeLa-*Anxa2^−/−^* cells infected with EMCV, VSV or HSV-1 were also higher than that of in the WT HeLa cells (Fig S3D-L). Moreover, the VSV and EMCV genomic RNA copy number and the HSV-1 genomic DNA copy number were significantly lower in the HeLa-*Anxa2^−/−^* cells than that of in the HeLa cells (Fig S3M-O). Collectively, these results indicated that the ANXA2 deficiency enhanced the type I IFN production, which resulted in the inhibition of viral replication.

### ANXA2 deficiency enhances host antiviral responses *in vivo*

To further define the function of ANXA2 in inhibiting type І IFN production and host antiviral responses *in vivo*, *Anxa2^−/−^* mice and their WT littermates were challenged with EMCV via intraperitoneal injections. We found that the mRNA levels of *Ifnβ1* in the heart and brain from *Anxa2^−/−^* mice were significantly higher than those from WT mice after infection with EMCV for 48 h (Fig 3A and 3B) and 72 h (Fig 3C and 3D). Correspondingly, the protein levels of IFN-β in serum from *Anxa2^−/−^* mice were also significantly increased (Fig 3E). Consistent with these results, the EMCV genomic RNA copy number in the heart were significantly lower in *Anxa2^−/−^* mice than that of their WT littermates (Fig 3F). Fewer signs of severe inflammation were observed in the brain and heart tissues from *Anxa2^−/−^* mice than in those from their WT littermates (Fig 3G). Additionally, brain tissues from WT littermates infected with EMCV exhibited severe glial cell and nerve cell degenerative necrosis, while those from *Anxa2^−/−^* mice exhibited few signs of glial cell and nerve cell degeneration (Fig 3G and 3H).

**Figure 3.**
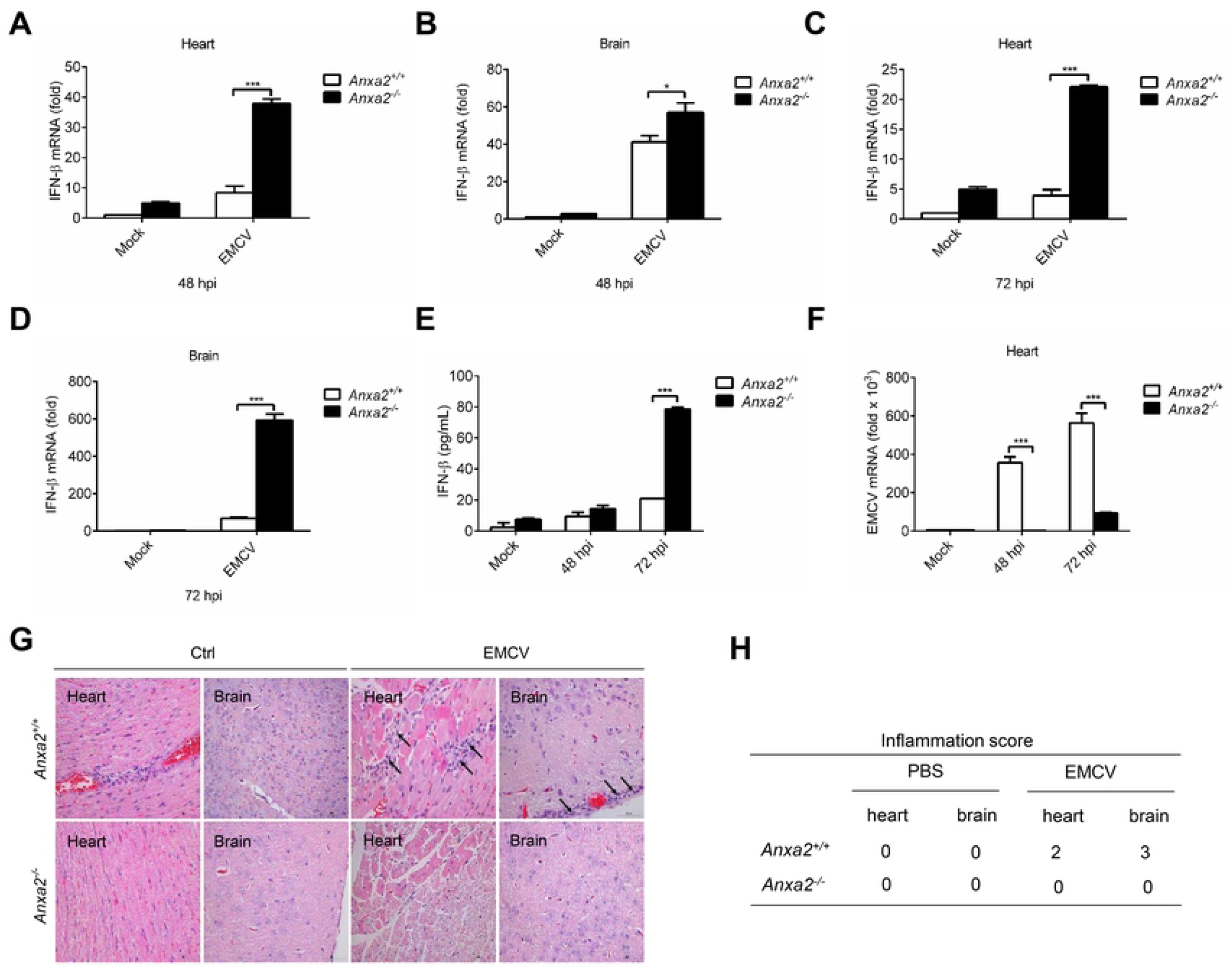
ANXA2 deficiency positively regulates antiviral responses in vivo. (A-D) The *Anxa2^+/+^* and *Anxa2^−/−^* mice (four mice per group) infected by the intraperitoneal injection of EMCV (2 × 10^5^ plaque-forming units (PFU) per mouse) for 48 h (A and B) and 72 h (C and D). The mRNA levels of *Ifnβ1* in the heart (A, C) and brain (B, D) were analyzed by qPCR analysis. The mRNA results are presented relative to those of mock infected WT cells. (E) Detection of the IFN-β levels in serum from mice by ELISA as in A and C. (F) qPCR analysis of EMCV RNA in the heart of *Anxa2^+/+^* and *Anxa2^−/−^* mice as in A and C; results are presented as in A and C. (G) Hematoxylin and eosin-stained images of heart and brain sections from *Anxa2^+/+^* and *Anxa2^−/−^* mice infected with EMCV for 96 h. Scale bars, 50 μm. (H) Inflammation score of (G): normal = 0, mild = 1, moderate = 2, and severe = 3 (n≥3, average score). \**P* < 0.05, \*\**P* < 0.01 and \*\*\**P* < 0.001 (two-tailed Student’s t-test (A-G)). Data are representative of three independent experiments with three biological replicates (mean ± s.d. in A-F) or are representative of three independent experiments with similar results (G).

### ANXA2 inhibits type I IFN production upstream of IRF3 phosphorylation

To elucidate the underlying molecular mechanisms by which ANXA2 negatively regulates type І IFN production, we first assessed the effect of ANXA2 on the IFN-β promoter activation induced by key molecules in the type І IFN signal pathway in HEK293T cells. As shown in Fig S4A-F, ectopically expressed ANXA2 significantly decreased the IFN-β promoter activation induced by RIG-I, MDA5, cGAS+STING, MAVS, TBK1, or IKKε in a dose-dependent manner, while the activation of IFN-β promoter induced by IRF3-5D (a constitutively active IRF3) was not affected (Fig S4G). To further confirm these results, HEK293T cells were transfected with a plasmid expressing RIG-I, MDA5, cGAS+STING, MAVS, TBK1, or IKKε, along with HA-ANXA2, the qPCR analysis showed that the ectopic expression of ANXA2 reduced the mRNA levels of *Ifnβ1* induced by the indicated molecules, but not by IRF3-5D (Fig 4A). Subsequently, HEK293T-*Anxa2^+/+^* and HEK293T-*Anxa2^−/−^* cells were transfected with a plasmid expressing RIG-I, MDA5, cGAS+STING, MAVS, TBK1, or IKKε, along with IFN-β and ISRE promoter reporters. We found that the IFN-β and ISRE promoter activation mediated by RIG-I, MDA5, cGAS+STING, MAVS, TBK1 and IKKε was significantly increased in HEK293T-*Anxa2^−/−^* cells compared to HEK293T-*Anxa2^+/+^* cells. Additionally, overexpression of ANXA2 in HEK293T-*Anxa2^−/−^* cells restored the inhibitory effects of ANXA2 on RIG-I-, MDA5-, cGAS+STING-, MAVS-, TBK1- and IKKε-mediated IFN-β and ISRE promoter activation but not IRF3-5D (Fig 4B, C). These results suggest that ANXA2 may inhibit type І IFN production upstream of IRF3 phosphorylation.

**Figure 4.**
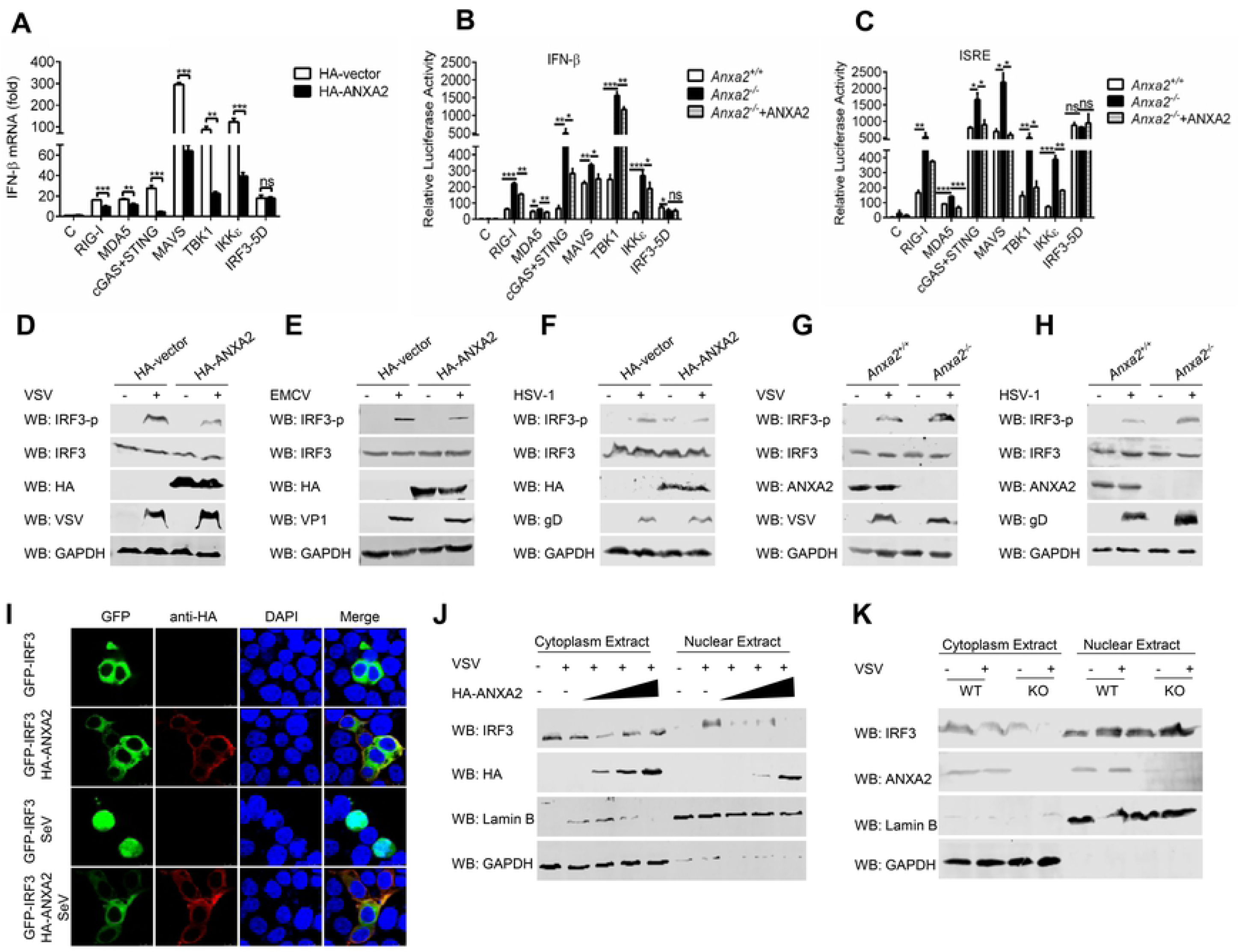
ANXA2 inhibits type I IFN production upstream IRF3 phosphorylation. (A) qPCR analysis of the mRNA levels of *Ifnβ1* in the HEK293T cells transfected with a plasmid expressing RIG-I, MDA-5, MAVS, cGAS+STING, TBK1, IKKε or IRF3-5D, along with an empty vector or a plasmid expressing ANXA2. (B and C) Luc activity of the IFN-β-luc (A) or ISRE-luc (B) reporter in the HEK293T-*Anxa2^+/+^* and HEK293T-*Anxa2^−/−^* cells transfected with an IFN-β-luc or ISRE-Luc reporter and a Renilla-TK reporter together with a plasmid expressing RIG-I, MDA-5, MAVS, cGAS+STING, TBK1, IKKε or IRF3-5D, along with an empty vector or a plasmid expressing ANXA2. (D-F) HEK293T cells were transfected with an empty vector or a plasmid expressing HA-ANXA2 and then mock infected or infected with VSV (D), EMCV (E) or HSV-1 (F). Immunoblot analysis of IRF3, phosphorylated IRF3, EMCV VP1 protein, HSV-1 gD protein, and GAPDH. (G, H) HEK293T-*Anxa2^+/+^* and HEK293T-*Anxa2^−/−^* cells were mock infected or infected with VSV (G) or HSV-1 (H). The cells were collected for immunoblot analysis of IRF3, phosphorylated IRF3, EMCV protein, HSV-1 protein, and GAPDH. (I) HeLa cells were transfected with a plasmid encoding ANXA2 or IRF3 alone or both. At 24 hpt, the cells were then infected with SeV for 12 h. The localization of IRF3 was detected by laser scanning confocal microscope. (J) HEK293T cells were transfected with increasing a plasmid encoding ANXA2, and then infected with VSV. IRF3 in the nuclear and cytoplasmic compartments was detected by Western blotting. (K) HEK293T-*Anxa2^+/+^* and HEK293T-*Anxa2^−/−^* cells were infected with VSV. IRF3 in the nuclear and cytoplasmic compartments was detected by Western blotting. Lamin B and GAPDH were used as nuclear and cytosolic markers.

To examine whether ANXA2 inhibits IRF3 phosphorylation, HEK293T cells were transfected with an empty vector or a plasmid expressing HA-tagged ANXA2, respectively. These cells were then mock infected or infected with VSV, EMCV or HSV-1. We found that ectopically expressed ANXA2 significantly inhibited the phosphorylation of IRF3 induced by VSV, EMCV or HSV-1 infection (Fig 4D-F). Similarly, ectopically expressed ANXA2 reduced poly(I:C)- or SeV-induced IRF3 phosphorylation in a dose-dependent manner (Fig S5A and 5B). Subsequently, primary peritoneal macrophages from *ANXA2^−/−^* mice and their wildtype littermates were infected with VSV or HSV-1. As shown in Fig 4G and 4H, the phosphorylation levels of IRF3 in primary peritoneal macrophages from *ANXA2^−/−^* mice were higher than those from WT mice infected with VSV or HSV-1, although the total protein levels of IRF3 were not affected. Consistent with these results, the phosphorylation level of IRF3 in the HeLa-*Anxa2^−/−^* cells was higher than that in the wild type HeLa cells, and overexpression of ANXA2 in HeLa-*Anxa2^−/−^* cells restored the inhibitory effects of ANXA2 on SeV-mediated phosphorylation of IRF3 (Fig S5C).

It has been shown that the phosphorylation of IRF3 results in its translocation to nucleus. Therefore, we further studied the effect of ANXA2 on IRF3 nuclear translocation. As shown in Fig 4I and Fig S5D, the amount of nuclear-translocated IRF3 was significantly reduced following ANXA2 expression upon SeV and EMCV infection. In agreement with these results, Western blot analysis of the protein level of IRF3 in the cytoplasmic and nuclear fractions showed that the amount of nuclear translocation of IRF3 induced by VSV infection decreased in a dose-dependent manner with the increasing expression of ANXA2, while knockout of ANXA2 expression significantly enhanced the nuclear translocation of IRF3 upon VSV infection (Fig 4J and 4K). Overall, our findings reveal that ANXA2 inhibits the phosphorylation and nuclear translocation of IRF3 induced by DNA virus or RNA virus infection.

### ANXA2 inhibits the interaction of MDA5-MAVS and MAVS-TRAF3 induced by RNA virus

To identify the target of ANXA2, we examined the interaction between ANXA2 and key molecules involved in RLRs-mediated and cGAS-STING signaling pathway including RIG-I, MDA5, cGAS, STING, MAVS, TBK1, IKKε and IRF3. As shown in Fig 5A, ANXA2 co-immunoprecipitated with MDA5, MAVS, STING and IRF3 when these proteins were co-expressed with these proteins in HEK293T cells. To validate the interaction between ANXA2 and MDA5, plasmids expressing HA-ANXA2 and Flag-MDA5 were co-transfected into HEK293T cells. The results showed that ANXA2 and MDA5 were co-immunoprecipitated (Fig 5B). Consistent with the result, endogenous MDA5 also interacted with endogenous ANXA2 regardless of mock infection or infection of EMCV (Fig 5C). To identify which domain of MDA5 is necessary for its interaction with ANXA2, five truncated mutants of MDA5 (MDA5-ΔC1, MDA5-ΔC2, MDA5-ΔATP, MDA5-ΔCTD and MDA5-CARD) were constructed, and Co-IP experiments were performed. As shown in Fig 5D, ANXA2 interacted with MDA5-WT, MDA5-ΔC1, MDA5-ΔATP, MDA5-ΔCTD and MDA5-CARD but not with MDA5-ΔC2, suggesting that the CARD domain of MDA5 is required for its interaction with ANXA2. Upon EMCV infection, MDA5 firstly senses EMCV genomic RNA, and then recruits MAVS through its CARD domain[28]. Therefore, we tested the interaction between MDA5 and MAVS and found that ANXA2 inhibited the recruitment of MAVS by MDA5 (Fig 5E). Consistent with the result, the interaction between MDA5 and MAVS was increased in HEK293T-*Anxa2^−/−^* cells compared with the HEK293T-*Anxa2*^+/+^ cells upon EMCV infection (Fig 5F).

**Figure 5.**
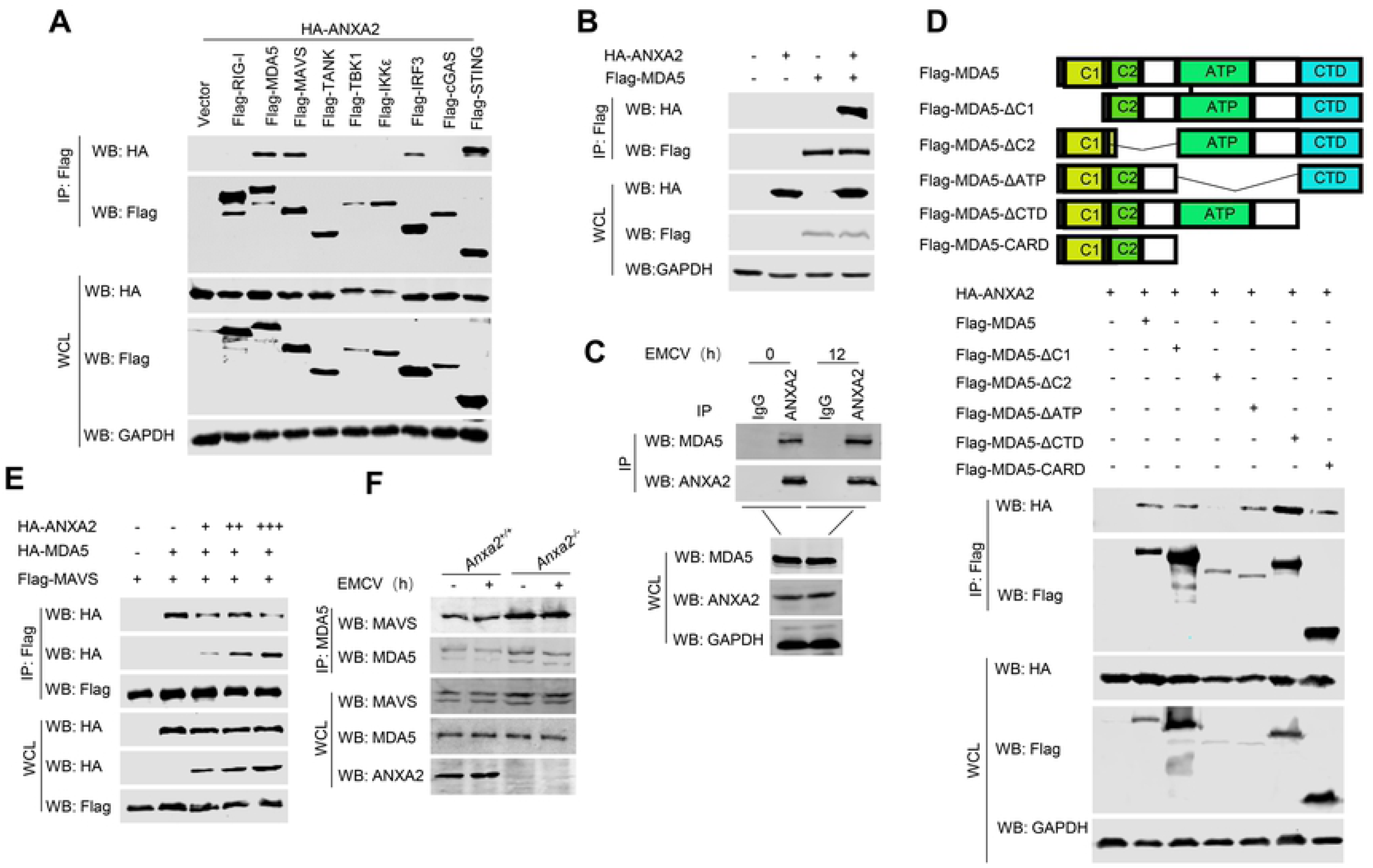
ANXA2 inhibits the recruiment of MAVS by MDA5. (A) Co-IP analysis of the interaction between ANXA2 and immune molecules in the HEK293T cells transfected with plasmids expressing HA-ANXA2 and Flag tagged RIG-I, MDA-5, MAVS, cGAS+STING, TBK1, IKKε or IRF3 as indicated. IP, immunoprecipitation. (B) Co-IP analysis of the interaction between ANXA2 and MDA5 in HEK293T cells transfected with plasmids expressing HA-ANXA2 and Flag-MDA5. (C) Co-IP analysis of the interaction between endogenous ANXA2 and MDA5 in mouse peritoneal macrophages that were mock infected or infected with EMCV. (D) MDA5 and its truncation mutants (top). Co-IP analysis of the interaction between ANXA2 and MDA5 or its deleted mutants in HEK293T cells transfected with a plasmid expressing HA-ANXA2 together with vector or Flag-MDA5 and its deleted mutants (below). (E) Co-IP analysis of the interactions among ANXA2, MDA5 and MAVS in the HEK293T cells transfected with plasmids expressing Flag-MAVS together with HA-MDA5 and vector or increasing amount of a plasmid encoding HA-ANXA2. (F) Co-IP analysis of the interaction between endogenous MDA5 and MAVS in mouse peritoneal macrophages that were mock infected or infected with EMCV.

The interaction between ectopic expressed ANXA2 and MAVS or between endogenous ANXA2 and MAVS in mock infected or EMCV infected HEK293T cells were examined by Co-IP (Fig 6A and 6B). Similarly, we generated four deletion mutants of MAVS bearing various combinations of the different domains to identify its binding domain to ANXA2. The results showed that MAVS-WT, MAVS-ΔCARD, MAVS-ΔN and MAVS-ΔTM interacted with ANXA2, but not MAVS-N, suggesting that ANXA2 interacted with MAVS is dependent on the linker between CARD and TM of MAVS (Fig 6C). It showed that the linker between CARD and TM of MAVS is necessary for its recruitment of TRAFs[29]. To detect the effect of ANXA2 on the interaction between MAVS and TRAF3, a plasmid expressing Flag-MAVS was transfected into HEK293T cells alone or together with a plasmid expressing HA-TRAF3 and increasing amount of a plasmid expressing GFP-ANXA2. As shown in Fig 6D, ANXA2 inhibited the interaction between MAVS and TRAF3 in a dose-dependent manner. Consistent with the above results, the interaction between MAVS and TRAF3 was enhanced in HEK293T-*Anxa2^−/−^* cells compared to that in the wile type HEK293T cells upon EMCV infection (Fig 6E). Of note, the location of MAVS on the mitochondria was not affected upon overexpression or deletion of ANXA2 (Fig 6F and Fig 6G).

**Figure 6.**
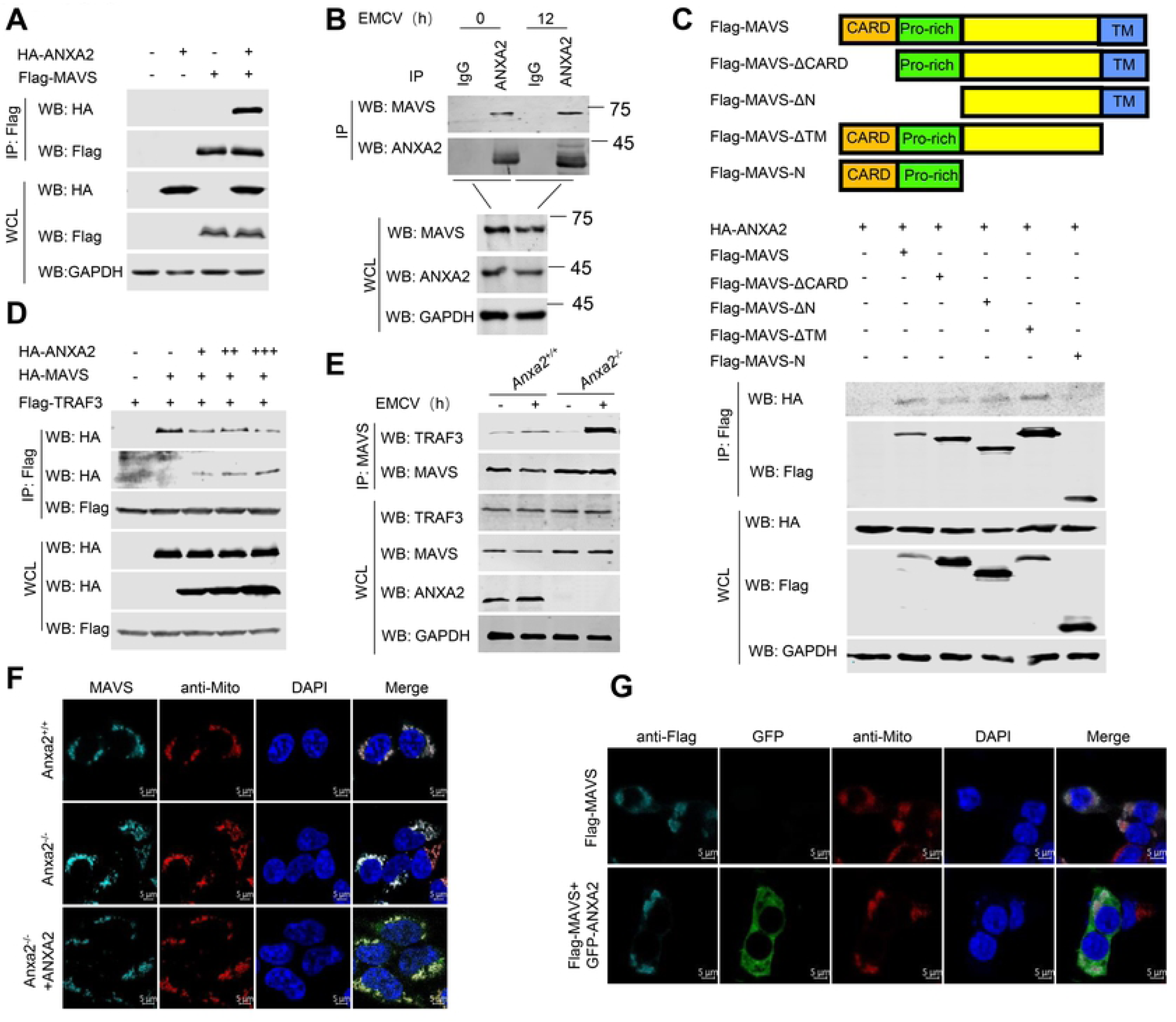
ANXA2 disrupts the interaction between MAVS and TRAF3. (A) Co-IP analysis of the interaction between ANXA2 and MAVS in HEK293T cells transfected with plasmids expressing HA-ANXA2 and Flag-MAVS. (B) Co-IP analysis of the interaction between endogenous ANXA2 and MAVS in mouse peritoneal macrophages that were mock infected or infected with EMCV. (C) MAVS and its truncation mutants (top) and Co-IP analysis of the interaction between ANXA2 and MAVS or its deleted mutants in HEK293T cells transfected with a plasmid expressing HA-ANXA2 together with vector or Flag-MAVS and its deleted mutants (below). (D) Co-IP analysis of the interactions among ANXA2, TRAF3 and MAVS in the HEK293T cells transfected with plasmids expressing Flag-MAVS together with HA-TRAF3 and vector or increasing mount of a plasmid encoding HA-ANXA2. (E) Co-IP analysis of the interaction between endogenous MAVS and TRAF3 in mouse peritoneal macrophages that were mock infected or infected with EMCV. (F) The subcellular localization of MAVS in HEK293T cells expressing Flag-MAVS alone or together with GFP-ANXA2 was detected by immunofluorescence microscopy. Scale bars, 5 μm. (G) The subcellular localization of MAVS in HEK293T-*Anxa2^+/+^* and HEK293T-*Anxa2^−/−^* cells were detected by immunofluorescence microscopy. Scale bars, 5 μm.

Taken together, these data demonstrate that ANXA2 not only inhibits the recruitment of MAVS by MDA5 through interaction with the CARD domain of MDA5, but also inhibits the recruitment of TRAF3 by MAVS through interaction with the linker between CARD and TM of MAVS.

### ANXA2 inhibits the localization of STING on Golgi apparatus induced by DNA virus

STING has been recognized as an activator of immune responses via TBK1/IRF3 pathway, and it is suggested to play critical roles in host defense against DNA virus[11, 30]. To confirm the interaction between ANXA2 and STING, plasmids expressing Flag-STING and HA-ANXA2 were co-transfected into HEK293T cells. The results of Co-IP showed that Flag-STING interacted with HA-ANXA2 (Fig 7A). In addition, we found that interactions existed between endogenous ANXA2 and endogenous STING in HEK293T cells with or without HSV-1 infection (Fig 7B). To map the STING domain required for the interaction with ANXA2, we constructed four different plasmids encoding Flag-tagged deleted STING mutants (STING-TM, STING-ΔC, STING-CDN, STING-ΔTM). The results revealed that ANXA2 interacted with STING-TM and STING-ΔC, but not STING-CDN or STING-ΔTM, suggesting that the TM of STING is dispensable for the interaction between ANXA2 and STING (Fig 7C). Next, we further explored whether ANXA2 affects the positioning of STING in the Golgi apparatus after activation. As shown in Fig 7D, the ectopic expression of cGAS promotes the transport of STING from the ER to the Golgi, whereas ANXA2 inhibits the localization of STING on the Golgi apparatus.

**Figure 7.**
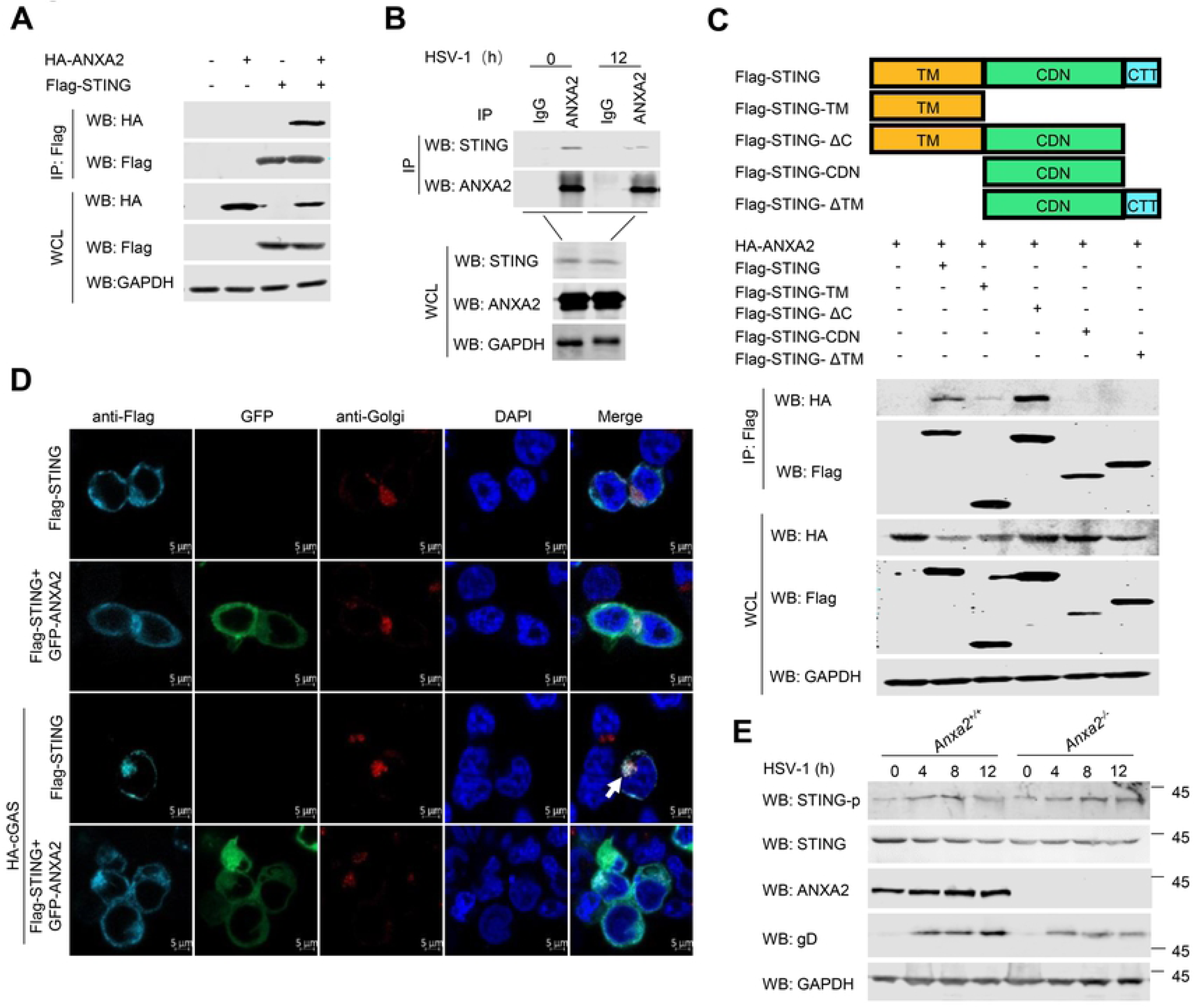
ANXA2 inhibits the location of STING on Golgi apparatus. (A) Co-IP analysis of the interaction between ANXA2 and STING in the HEK293T cells transfected with plasmids expressing HA-ANXA2 and Flag-STING. (B) Co-IP analysis of the interaction between endogenous ANXA2 and STING in mouse peritoneal macrophages that were mock infected or infected with HSV-1. (C) ANXA2 and its truncation mutants (top) and Co-IP analysis of the interaction between ANXA2 and STING or its deleted mutants in HEK293T cells transfected with a plasmid expressing Flag-STING together with vector or HA-ANXA2 and its deleted mutants. (D) The subcellular localization of STING in HEK293T cells expressing Flag-STING alone or together with GFP-ANXA2 as detected by immunofluorescence microscopy. Scale bars, 5 μm. (E) Western blotting analysis of STING phosphorylation upon HSV-1 infection mouse peritoneal macrophages for 0, 4, 8, 12 h.

### ANXA2 disrupts TBK1-IKKε-IRF3 complex formation mediated by both RNA virus and DNA virus

Both DNA and RNA virus activate TBK1/IRF3 pathways. To detect the interaction between ANXA2 and IRF3, plasmids expressing Flag-IRF3 and HA-ANXA2 were co-transfected into HEK293T cells. The result of Co-IP showed that Flag-IRF3 interacted with HA-ANXA2 (Fig 8A). In addition, we also found that interactions existed between endogenous ANXA2 and endogenous IRF3 in HEK293T cells infected with VSV (Fig 8B) or HSV-1 (Fig 8C). Immunofluorescence staining revealed that ANXA2 colocalized with IRF3 in the cytoplasm (Fig 8D). IRF3 contains four different domains: an N-terminal DNA binding domain (DBD), an IRF association domain (IAD), a C-terminal regulatory domain (RD), and a nuclear export sequence (NES). To map the regions of IRF3 responsible for the interaction with ANXA2, five different plasmids encoding Flag-tagged deleted IRF3 mutants (IRF3-D_1_: 1-382 aa, IRF3-D_2_: 1-357 aa, IRF3-D_3_: 1-140 aa, IRF3-D_4_: 141-427 aa, IRF3-D_5_: 201-427 aa) were constructed, and Co-IP experiments were performed. The results revealed that ANXA2 interacted with IRF3-D_1_, IRF3-D_2_ and IRF3-D_4_, but not IRF3-D_3_ or IRF3-D_5_, suggesting that the NES domain of IRF3 is required for its interaction with ANXA2 (Fig 8E). ANXA2 contains two different domains: a C-terminal domain with four repeats and a unique N-terminal domain[31, 32]. To map the ANXA2 domain required for the interaction with IRF3, we constructed six different plasmids encoding HA-tagged deleted ANXA2 mutants (ANXA2-D_1_: 1-102 aa, ANXA2-D_2_: 1-174 aa, ANXA2-D_3_: 1-259 aa, ANXA2-D_4_: 50-340 aa, ANXA2-D_5_: 122-340 aa, ANXA2-D_6_: 207-340 aa). As shown in Fig 8F, ANXA2-D_2_, ANXA2-D_3_, ANXA2-D_4_, ANXA2-D_5_ and ANXA2-D_6_ interacted with IRF3, but not ANXA2-D_1_, suggesting that the C-terminal region, especially the four repeated sequence, is indispensable for the interaction between ANXA2 and IRF3. Next, we tried to identify which domain of ANXA2 is required for its inhibition of TBK1-mediated IFN-β production. We found that ANXA2-D_2_, ANXA2-D_3_, ANXA2-D_4_ and ANXA2-D_5_, but not ANXA2-D_1_ or ANXA2-D_6_, inhibited IFN-β-luc reporter activation and the mRNA expression levels induced by TBK1 (Fig 8G and 8H).

**Figure 8.**
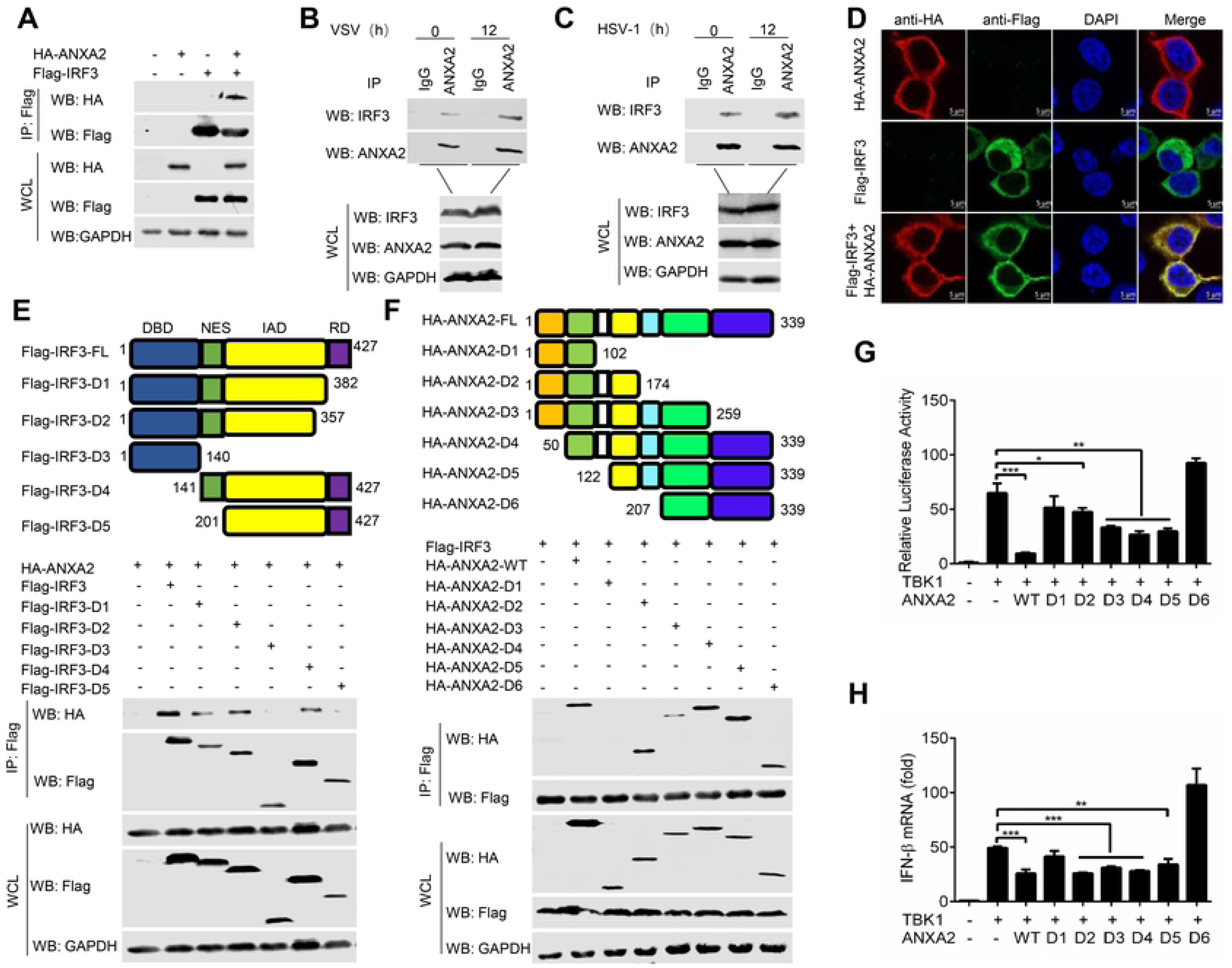
ANXA2 interacts with IRF3. (A) *In vitro* analysis of the interaction between ANXA2 and IRF3 in the HEK293T cells transfected with plasmids expressing HA-ANXA2 and Flag-IRF3. (B) Immunoprecipitation and immunoblot analysis of the interaction of endogenous ANXA2 and IRF3 in mouse peritoneal macrophages infected with VSV (B) or HSV-1 (C) at the indicated times. (D) The subcellular localization of ANXA2 and IRF3 in HEK293T cells expressing HA-ANXA2 and Flag-IRF3 as detected by immunofluorescence microscopy. Scale bars, 5 μm. (E) IRF3 and its deleted mutants (top) and the Co-IP analysis of the interaction between HA-ANXA2 and Flag-IRF3 or its deleted mutants in HEK293T cells (below). (F) Co-IP analysis of the interaction between Flag-IRF3 and HA-ANXA2 or its deleted mutants in HEK293T cells. (G) Luc activity of the IFN-β-luc reporter in HEK293T cells transfected with an IFN-β-Luc reporter and a Renilla-TK reporter, together with a plasmid expressing Flag-TBK1 and HA-ANXA2 or its deleted mutants. (H) qPCR analysis of *Ifnβ1* mRNA levels in HEK293T cells transfected with a plasmid expressing Flag-TBK1 and HA-ANXA2 or its mutants. \*\*\**P* < 0.001 (two-tailed Student’s t-test). Data are representative of three independent experiments with three biological replicates (mean ± s.d. in A-C).

Previous studies showed that TBK1 and IKKε are TRAF family member-associated NF-κB activator (TANK)-binding partners and the tricomplex (TBK1-IKKε-IRF3) is involved in participating in the type I IFN production[33, 34]. Activated TBK1 and IKKε phosphorylates the serine residues in IRF3, causing conformational changes and homogeneous dimerization of IRF3, followed by its nuclear translocation and activation of target gene transcription [35–37]. Therefore, we speculated that ANXA2 might inhibit IRF3 phosphorylation by blocking the interaction between TBK1 or IKKε and IRF3. The Co-IP results confirmed our hypothesis, as overexpressed ANXA2 markedly disrupted the interaction between TBK1 and IRF3 (Fig 9A) as well as the interaction between IKKε and IRF3 (Fig 9B) in a dose-dependent manner. Consistent with these results, the lack of ANXA2 expression significantly increased the interaction of the endogenous TBK1-IRF3 and IKKε-IRF3 upon VSV and HSV-1 infection in the HEK293T cells (Fig 9C and 9D) or mouse primary peritoneal macrophages (Fig 9E and 9F). Overall, our findings reveal that ANXA2 competes with TBK1 or IKKε in binding to the IRF3, leading to the inhibition of IRF3 phosphorylation and nuclear translocation.

**Figure 9.**
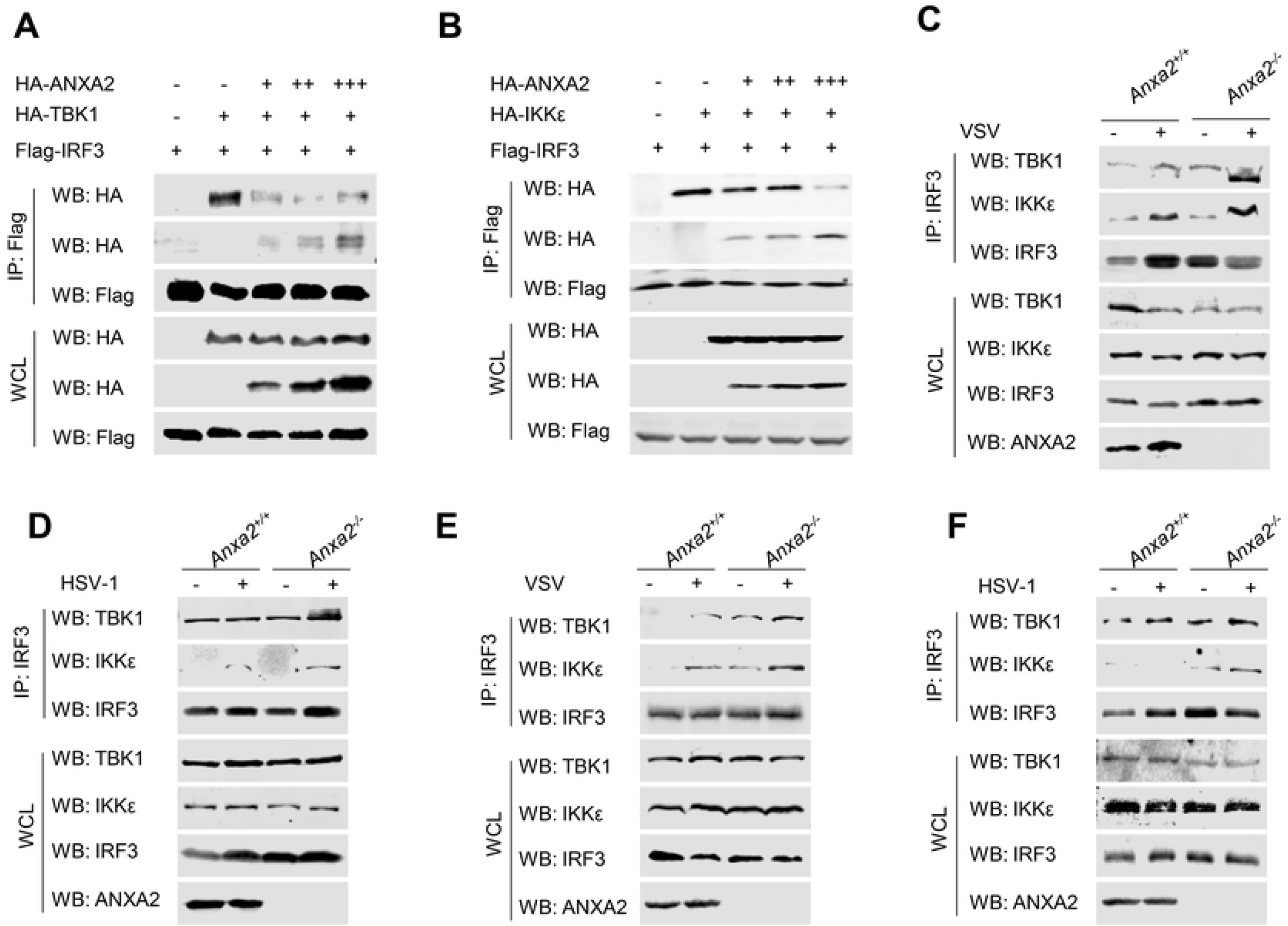
ANXA2 disrupts TBK1-IKKε-IRF3 complex. (A, B) HEK293T cells were transfected with a plasmid encoding Flag-IRF3 alone or together with a plasmid expressing HA-TBK1 (A) or HA-IKKε (B) and different amounts of plasmids encoding HA-ANXA2 as indicated. At 24 hpi, Co-IP was performed with anti-FLAG. (C-F) Immunoprecipitation and immunoblot analysis of the interaction of endogenous TBK1 or IKKε and IRF3 in HEK293T-*Anxa2^+/+^* and HEK293T-*Anxa2^−/−^* cells (C, D) or mouse peritoneal macrophages (E, F) mock infected or infected with VSV or HSV-1.

Taken together, both RNA virus and DNA virus infection induced ANXA2 expression to inhibit type I IFN production. Mechanistically, ANXA2 not only inhibits the recruitment of MAVS by MDA5 and the interaction between MAVS and TRAF3 upon RNA virus infection, but also inhibits the location of STING on Golgi apparatus upon DNA virus infection. Additionally, ANXA2 disrupts TBK1/IKKε/IRF3 complex formation through competing with TBK1 or IKKε binding to IRF3, which leads to reduction of phosphorylation and nuclear translocation of IRF3 and decreased type I IFN production (Fig 10).

**Figure 10.**
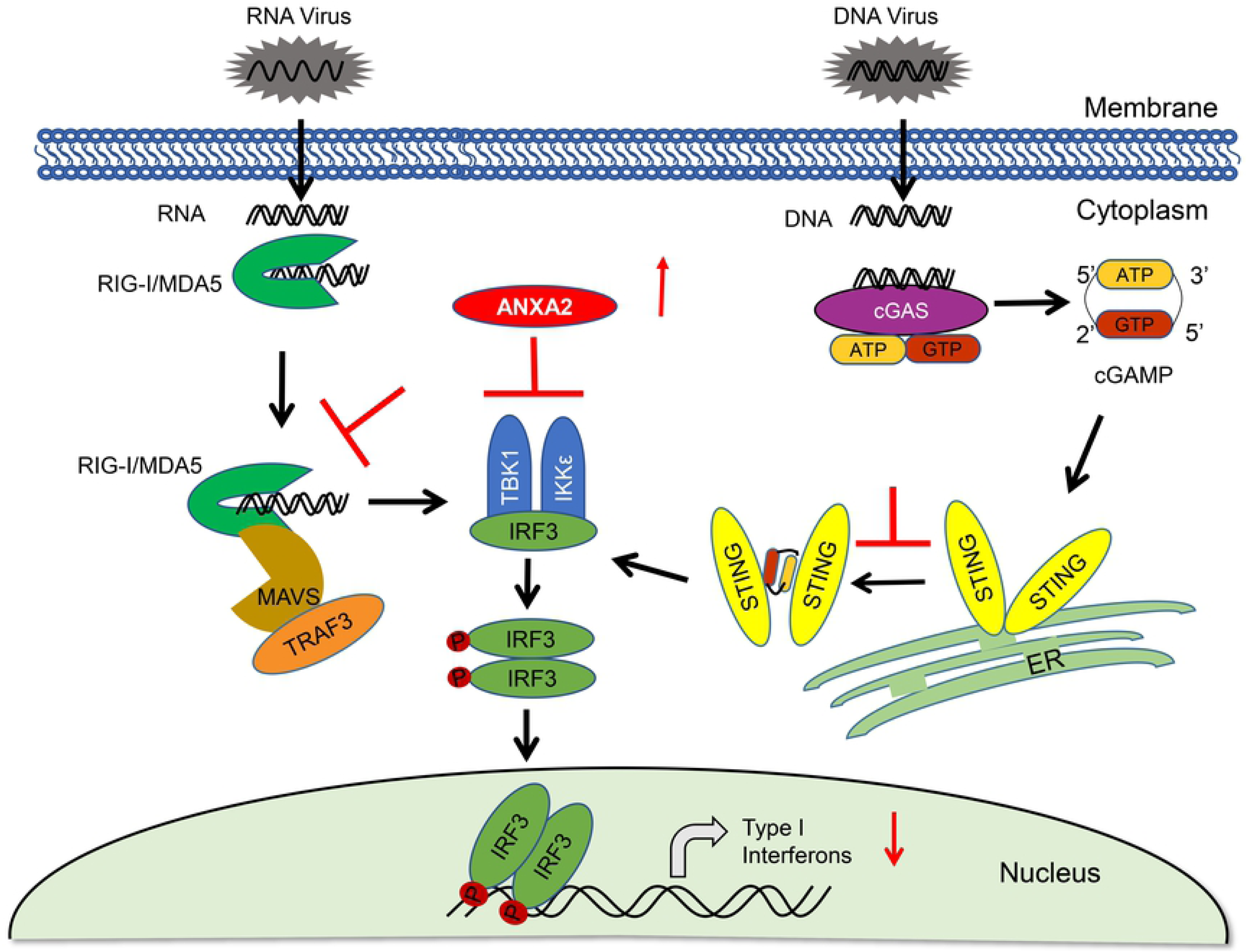
Schematic model of ANXA2’s inhibition of type I IFN production. Upon virus infection, viral genomic RNA was recognized by MDA5/RIG-I, and viral genomic DNA was recognized by cGAS to induce type I IFN production to activate host antiviral responses. To antagonize host antiviral innate immune response, both RNA virus and DNA virus infection induce ANXA2 expression, which not only inhibits the interaction between MDA5 and MAVS induced by RNA virus but also inhibits the localization of STING on Golgi apparatus induced by DNA virus. In addition, ANXA2 also competes with TBK1 or IKKε to bind to IRF3 to inhibit IRF3 phosphorylation and nuclear translocation.

## Discussion

Annexins are a family of evolutionary conserved multifunctional proteins, which widely distributed in various tissues and cells of plants and animals, and can reversibly bind to phospholipid membranes and to calcium ions (Ca^2+^)[17]. Annexins can not only participate in the occurrence of various membrane events regulated by Ca^2+^, such as membrane transport, cytoskeleton formation[38, 39], and ion channel establishment[40–42], but also are related to cell apoptosis and tumorigenesis[43–45]. Recently, increasing evidence has demonstrated that annexins play important roles in innate immune and inflammatory responses[46, 47]. It has been reported that in the TLR4/TLR3-TRIF signaling pathway, the C-terminus of ANXA1 directly associates with TBK1 to promote the TLRs-mediated IFN-β production, indicating that ANXA1 plays an important role in regulating host antiviral responses[48]. ANXA1 is not only related to IFN production signaling pathways, but can also affect IFN downstream signaling pathway. For example, ANXA1 rapidly promotes IFN-β production and IRF3 activation after RIG-I stimulation, while knockdown of ANXA1 expression delays the phosphorylation of IRF3 and STAT1, leading to lower expression of ISGs, such as IFIT1[49]. ANXA7, another Annexin member, enhances the IFN-β promoter activity induced by chicken MDA5 (chMDA5), thereby inhibiting the infection of the recombinant H5N1 virus (rNS1-SD30) lacking the eIF4GI binding domain of NS1[50].

As a membrane scaffold and binding protein, ANXA2 is mainly expressed in the plasma membrane and intracellular vesicles[18, 19], where it participates in the replication process of a variety of viruses. For example, ANXA2 interacts with the non-structural protein 1 (NS1) of AIV to increase the viral replication[26]. Our previous studies also showed that ANXA2 interacts with the non-structural proteins 9 (NSP9) of PRRSV to promote viral replication, and Vimentin interacts with the N protein of PRRSV in the case of ANXA2[27]. In this study, we noticed that virus infection induced ANXA2 expression, subsequently, we found that overexpression of ANXA2 inhibited both RNA viruses (SeV, VSV, or EMCV) and DNA virus (HSV-1)-induced IFN-β production, whereas ANXA2 deficiency enhanced type I IFN production and suppressed virus replication *in vitro* and *in vivo*. ANXA2 interacted with IRF3 to inhibit type I IFN production by blocking the phosphorylation and nuclear translocation of IRF3 induced by RNA virus and DNA virus.

RIG-I-like helicases, such as RIG-I, MDA5, and LGP2, act as important cytosolic PRRs to sense viral dsRNA. RIG-I and MDA5 transduces antiviral signal through interacting with MAVS via CARD domain. MAVS consists of three domains, including CARD domain, TM domain and the linker between CARD and TM. MAVS recruits TRAF3 through the linker between CARD and TM to activate downstream kinases such as TBK1 and IKKε to phosphorylate IRF3, leading to increased production of type I IFN and expression of antiviral genes[5, 51]. In this study, we found that ANXA2 not only interacted with MDA5 through CARD domain to inhibit its recruitment of MAVS but also interacted with the linker between CARD and TM of MAVS to inhibit the interaction between MAVS and TRAF3. Upon DNA virus infection, the cGAS senses and binds to the viral DNA and catalyzes formation of 2’-3’-cGAMP, an atypical cyclic di-nucleotide second messenger that can be sensed by STING. STING translocates from ER to Golgi, leading to phosphorylation of STING. In this study, we found that ANXA2 interacted with the TM domain of STING and inhibited its localization on Golgiosome and phosphorylation. Activated STING recruited and activatedTBK1[52]. Therefore, the type I IFN signaling triggered by RNA virus and DNA virus converges on TBK1. Activated TBK1 phosphorylates IRF3 and promotes its dimerization and translocation into the nucleus, where it forms an active transcriptional complex that binds to IFN promoter and triggers the type I IFN genes transcription[53, 54].

Normally, IRF3 mainly exists in the cytoplasm and can shuttle between cytoplasm and nucleus. The NLS and NES in IRF3 are constitutively active, but nuclear export is normally dominant. Following viral infection, activated TBK1 and IKKε interact with the IAD of IRF3 and phosphorylate IRF3, which promotes IRF3 to translocate into the nucleus to induce type I IFN production[15, 55]. IRF3 accumulates in the nucleus, and this accumulation relies on the function of its NLS[56]. To date, several studies demonstrated that some host proteins inhibit IRF3 activation by directly interacting with IRF3. For instances, TRIM26 binds to IRF3 and promotes its K48-linked polyubiquitination and degradation in nucleus[57]. Our previous study also showed that DDX19 inhibits TBK1- and IKKε-mediated phosphorylation of IRF3 by competing with TBK1 or IKKε binding to the IAD domain of IRF3[58]. In this study, we found that ANXA2 inhibited IRF3 phosphorylation and nuclear entry induced by both RNA virus and DNA virus by inhibiting TBK1/IKKε/IRF3 complex formation. Unexpectedly, ANXA2 competed with IRF3 to bind to TBK1/IKKε through interacting with the NES domain but not the IAD domain of IRF3. This may be caused by the steric hindrance related to the structure of ANXA2, which requires further study to confirm.

ANXA2 is an abundant protein that associates with biological membranes as well as the actin cytoskeleton. It has been implicated in intracellular vesicle fusion, the organization of membrane domains, lipid rafts and membrane-cytoskeleton contacts. It consists of a highly conserved core domain of four homologous repeats of 70–80 amino acids called the annexin repeats and a unique 30-amino-acid long N-terminal ‘head domain’[32, 59]. In this study, we found that 102-207 aa of ANXA2 is required to inhibit TBK1 induced IFN-β promoter activation, and 102-339 aa of ANXA2 is necessary to interact with IRF3.

ANXA2 plays an essential role in the regulation of innate immune response. A report demonstrated that ANXA2 plays an anti-inflammatory role in response to injury or viral infection[60]. Another previous study showed that ANXA2 has a role in limiting inflammation by promoting anti-inflammatory signals[61]. In agree with these, the mice lacking ANXA2 showed a lower survival rate when they were infected with bacteria, reflecting a dysregulated inflammatory response[62]. Using the ANXA2 knockout mice as model, we demonstrated that IFN-β production significantly increase in primary peritoneal macrophages from *Anxa2^−/−^* mice upon EMCV infection compared to that from wild type mice, which in turn inhibited viral replication. These data suggest that ANXA2 is a negative regulator of IFN-β production and antiviral immune response during viral infection *in vivo*. Therefore, the virus infection may limit IRF3 activation and IFN-β production by inducing ANXA2 expression, thereby promoting its escape from the host innate immune responses.

In summary, we identified a novel function of ANXA2 involved in host antiviral responses. Mechanistically, ANXA2 negatively regulates IFN-β production by targeting MDA5, MAVS, STING and IRF3. ANXA2 disrupts the interaction of MDA5-MAVS and MAVS-TRAF3 upon RNA virus infection while inhibits the localization of SITNG on Golgi apparatus upon DNA virus infection. ANXA2 also inhibits TBK1 and IKKε in binding to IRF3 through interaction with IRF3. Understanding these processes may shed light on ANXA2’s new function in viral infection-mediated type I IFN production. Therefore, this study provides a novel target for anti-viral drug design to prevent viral invasion.

## Materials and Methods

### Mice

*Anxa2^−/−^* mice generated by homologous recombination technology were purchaced from Saiye, Biotechnology Co., Ltd (Guangzhou, China). The mouse genotype was confirmed by PCR using the following primers: forward 5’-CAACTGAGGCACACTCACAAGCG-3’, and reverse 5′-GAGAAGGGCTGGCTTAGGGCACT-3′ and 5′-ACTGTGCTGTGAATGCCCACCTTG-3′. All mice were generated and housed in specific pathogen-free (SPF) barrier facilities at the Harbin Veterinary Research Institute (HVRI) of the Chinese Academy of Agricultural Sciences (CAAS) (Harbin, China). All animal experiments were performed according to animal protocols approved by the Subcommittee on Research Animal Care at the HVRI. Male and female *Anxa2^−/−^* and wild-type littermates (6-8 weeks old) were used throughout the experiments.

### Cell lines

Human HEK293T and HeLa cells were purchased from American Type Culture Collection (Manassas, VA). Peritoneal macrophages were isolated from mice 3 days after injection of thioglycollate (MERCK) and cultured in RPMI 1640 medium supplemented with 10% fetal bovine serum (FBS), 100 U/mL penicillin and 100 mg/mL streptomycin at 37°C with 5% CO_2_. Human HEK293T and HeLa cells were cultured in Dulbecco’s Modified Eagle’s Medium (DMEM) supplemented with 10% FBS, 100 U/mL penicillin, and 100 mg/mL streptomycin at 37°C with 5% CO_2_. *Anxa2^−/−^* and wild type HeLa cell lines were purchased from Edigene Inc. (Beijing, China).

### Viruses

The Sendai/Cantell strain (SeV strain, product code VR-907) was purchased from the American Type Culture Collection (ATCC) and amplified in specific pathogen-free eggs. The EMCV HB10 strain was provided by Prof. Shangjin Cui and the VSV and VSV-GFP were kindly provided by Prof. Zhigao Bu (HVRI, China). HSV-1 was kindly provided by Prof. Hongbin Shu (Wuhan University, China).

### Plasmids

Plasmids expressing Flag-tagged RIG-I, MDA5, cGAS, STING, MAVS, TANK, TBK1, IKKε, IRF3 and IRF3-5D have been previously described (Huang et al., 2015). The IFN-β reporter, ISRE reporter, and TK-Renilla reporter were obtained from Prof. Hong Tang. To construct plasmids expressing HA- or Flag-tagged ANXA2, cDNAs corresponding to the human ANXA2 gene were amplified by standard reverse transcription-polymerase chain reaction (RT-PCR) using total RNA extracted from HeLa cells as templates and were then cloned into the pCAGGS-HA/Flag vector. All constructs were validated by DNA sequencing. The primers used in this study are available upon request (Table 1).

### Viral infection

For qRT-PCR or immunoblot analysis, cells (2 × 10^5^) were plated 24 h before infection with various viruses at the indicated time points. For viral replication assays, peritoneal macrophages were infected with EMCV, VSV or HSV-1 for 0, 4, 8, 12 h. Virial replication was analyzed by qRT-PCR analysis. For mouse infection, six- to eight-week-old age- and sex-matched *Anxa2^+/+^* and *Anxa2^−/−^* littermates were intraperitoneally injected with EMCV (2 × 10^4^ PFU per mouse) or HSV-1 (2 × 10^7^ PFU per mouse). For survival experiments, the survival of animals was monitored every day after EMCV infection. The sera from EMCV-infected mice were collected for ELISA analysis at 48 h and 72 h post infection, and the heart and brain were collected for qRT-PCR, the EMCV titers or histological analysis.

### Luciferase reporter gene assay

Luciferase activities were measured with Dual-Luciferase Reporter Assay System (Promega), according to the manufacturer’s instructions. Data were normalized for transfection efficiency by the division of Firefly luciferase activity with that of Renilla luciferase.

### RNA extraction and qPCR

Total RNA was extracted using TRIzol reagent (Invitrogen), and the reverse transcription products were amplified using the Agilent-Strata gene MxReal-Time qPCR system with a PrimeScript^™^ RT Reagent Kit (Takara). Reverse transcription products were amplified using an Agilent-Strata gene MxReal-Time qPCR system with TB Green^®^ Premix Ex Taq^™^ II (Tli RNaseH Plus) (Takara) according to the manufacturer’s instructions. Data were normalized to the level of β-actin expression in each individual sample. The qPCR primers are listed in Table 2.

### IFN sensitivity assay

The cellular supernatants were collected and used to assess for their ability to inhibit vesicular stomatitis virus-GFP (VSV-GFP) replication. Briefly, the HEK293T cells, HEK293T-*Anxa2^−/−^* cells or HEK293T-*Anxa2^−/−^* cells overexpressing ANXA2 were infected with SeV at an MOI of 10, and the cell supernatants were collected and UV-deactivated using a 254 nm UV light source for 15 min. UV-deactivated cell supernatants were then diluted 1:10 in RPMI-1640 media and added to MDBK cells. Following a 24 h pre-treatment, MDBK cells were infected with VSV-expressing GFP at an MOI of 5.0 for a duration of 5-8 h. VSV infection was determined by fluorescence under a UV light source.

### Co-immunoprecipitation and Western blot analysis

Co-immunoprecipitation and Western blot analysis were performed as previously described [58]. Briefly, for Co-IP, whole cell extracts were lysed in lysis buffer (50 mM Tris-HCl, pH 7.4, 150 mM NaCl, 5 mM MgCl_2_, 1 mM EDTA, 1% Triton X-100, and 10% glycerol) containing 1 mM PMSF and 1 × protease inhibitor cocktail (Roche). Then, cell lysates were incubated with anti-Flag (M2) beads or with protein G Plus-Agarose immunoprecipitation reagent (Santa Cruz Biotechnology) together with 1 μg of the corresponding antibodies at 4°C overnight on a roller. The precipitated beads were washed five times with cell lysis buffer. For Western blot analysis, equal amounts of cell lysates and immunoprecipitants were resolved by 10-12% sodium dodecyl sulfate polyacrylamide gel electrophoresis (SDS-PAGE) and were then transferred to a polyvinylidene difluoride (PVDF) membrane (Millipore). After incubation with primary and secondary antibodies, the membranes were visualized by ECL chemiluminescence (Thermo Fisher Scientific) or an Odyssey two-color infrared fluorescence imaging system (LI-COR).

### Confocal microscopy analysis

HeLa cells or HEK293T cells were transfected with the indicated plasmids and were then fixed for 20 min in 4% paraformaldehyde in 1 × phosphate-buffered saline (PBS) at pH 7.4. Fixed cells were permeabilized for 20 min with 0.3% Triton X-100 in 1 × PBS and were then blocked in 1 × PBS with 10% bovine serum albumin (BSA) for 30 min. The cells were incubated with the appropriate primary antibodies and were then stained with Alexa Fluor 594-labeled goat anti-mouse immunoglobulin G and Alexa Fluor 488-labeled goat anti-rabbit IgG. The subcellular localization of indicated proteins were visualized using a Zeiss LSM-880 laser scanning fluorescence microscope (Carl Zeiss AG, Oberkochen, Germany) under a 63 × oil objective.

### Generation of *Anxa2* deficient mice

CRISPR/Cas9 genomic editing for gene deletion was used as previously described[63]. To create mammalian *Anxa2^−/−^* cells, one CRISPR guide RNA (sgRNA) sequence targeting the ANXA2 locus in the genome was chosen based on the specificity scores (http://crispr.mit.edu/). The sgRNA sequence that was used as follows: ANXA2 sgRNA, 5’-GCACTGAAGTCAGCCTTATCTGG-3’. The sgRNA sequence was cloned into the pSpCas9 (BB)-2A-GFP plasmid (pX458, Addgene). The construct was then independently transfected into HEK293T cells. Cells expressing GFP were isolated by flow cytometry, and single cells were seeded into the 96-well plates. After clonal expansion, ANXA2 protein expressions in different clones were analyzed by immunoblot, and the genomic DNAs from those clones that have undetectable ANXA2 protein expressions were extracted and amplified by PCR for ANXA2 gene sequencing.

### Histopathology analysis

The brains and hearts of WT and *Anxa2^−/−^* mice infected with the EMCV HB10 strain were fixed in 10% formalin neutral buffer solution overnight. The tissues were embedded in paraffin blocks, and then sectioned at a 4-μm thickness for staining with hematoxylin and eosin in accordance with standard procedures. The results were analyzed by light microscopy. Representative views of the brain and heart sections are shown.

### Statistical analysis

Statistical analysis was conducted using an unpaired Student’s t-test, a two-tailed Student’s t-test and one-way or two-way ANOVA followed by the Bonferroni post-test. P values less than 0.05 were considered statistically significant. For mouse survival studies, Kaplan-Meier survival curves were generated and analyzed for statistical significance with GraphPad Prism 6.0. Sample sizes were chosen by standard methods to ensure adequate power, and no exclusion, randomization of weight or sex or blinding was used for animal studies.

## Acknowledgments

This study was supported by the Natural Science Foundation of Heilongjiang Province of China (grants No. YQ2020C022), National Natural Science Foundation of China (grants No. 31941002) and the State Key Laboratory of Veterinary Biotechnology Program (grants No. SKLVBP202101).

---

**Figure S1.**
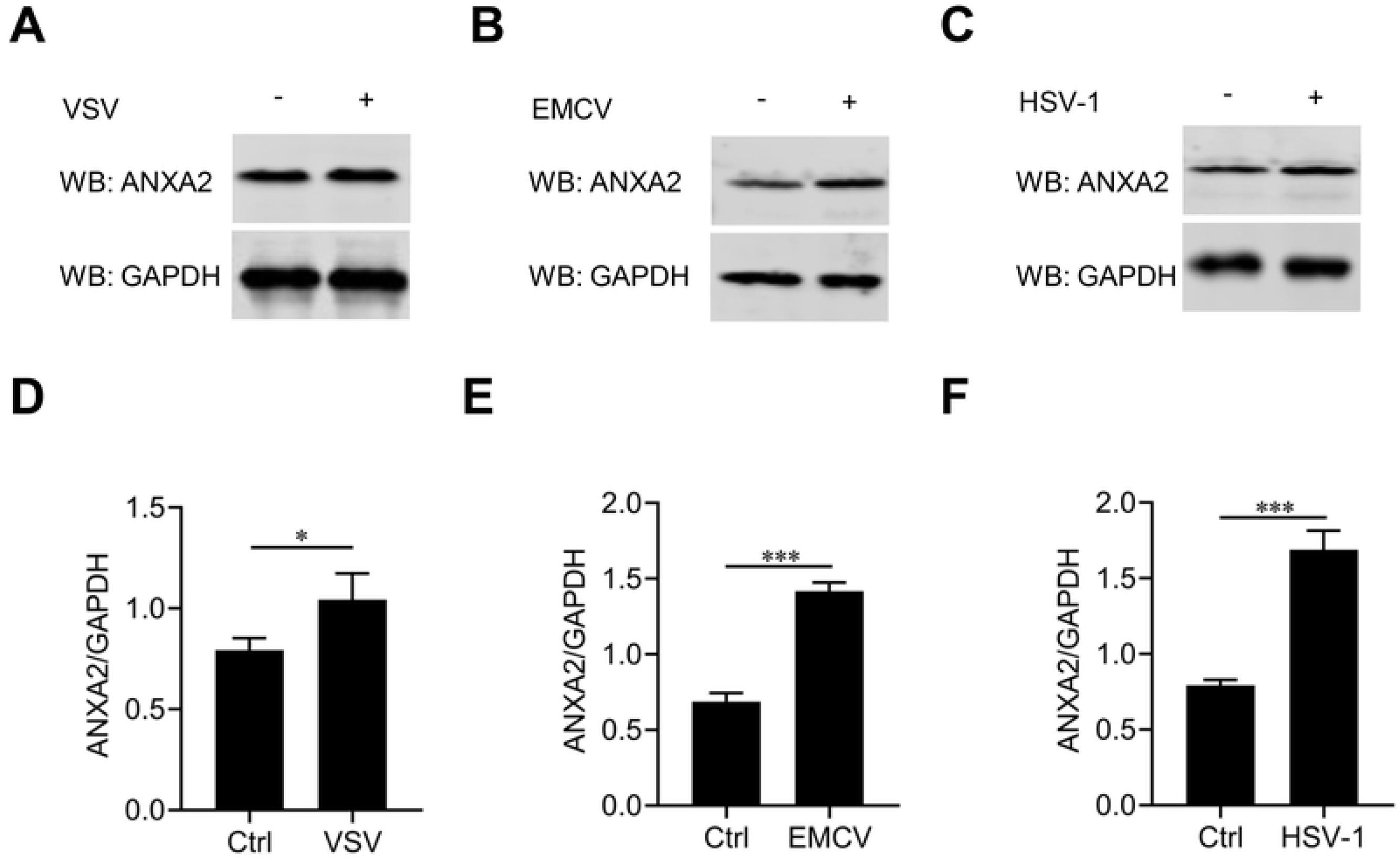

**Figure S2.**
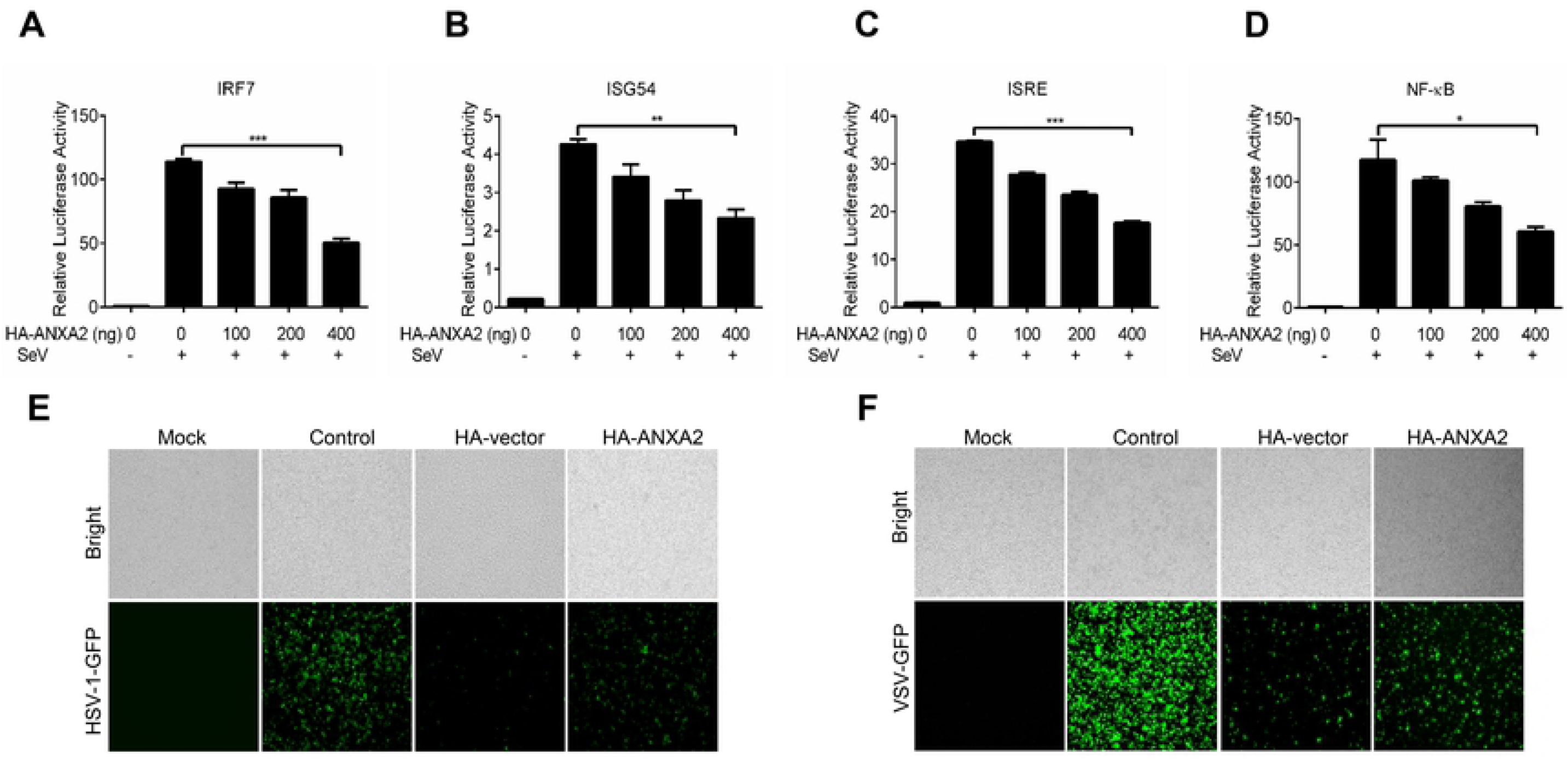

**Figure S3.**
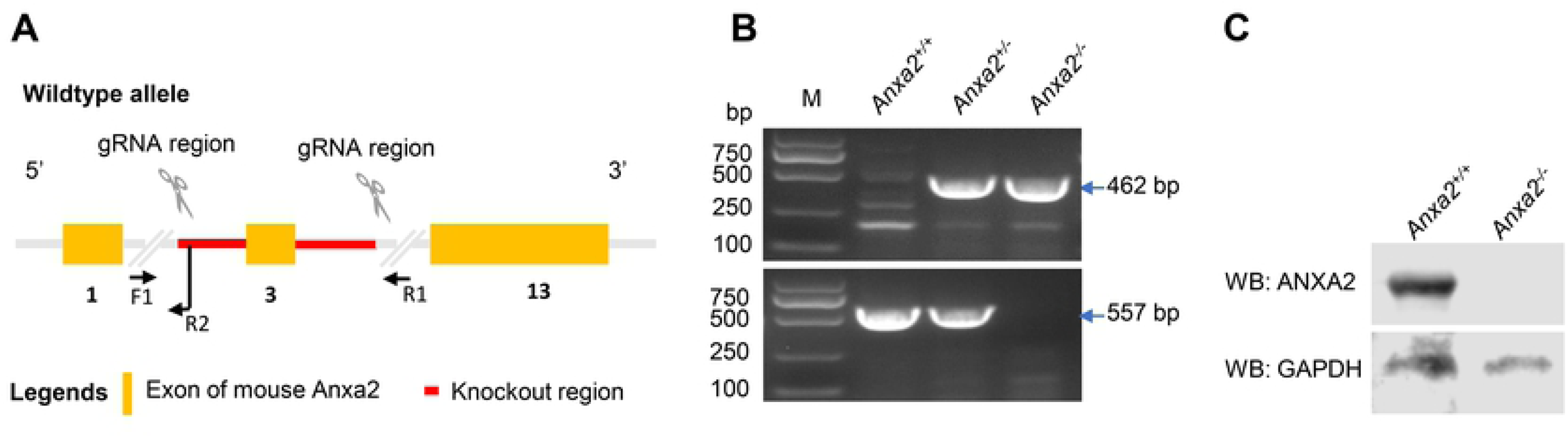

**Figure S4.**
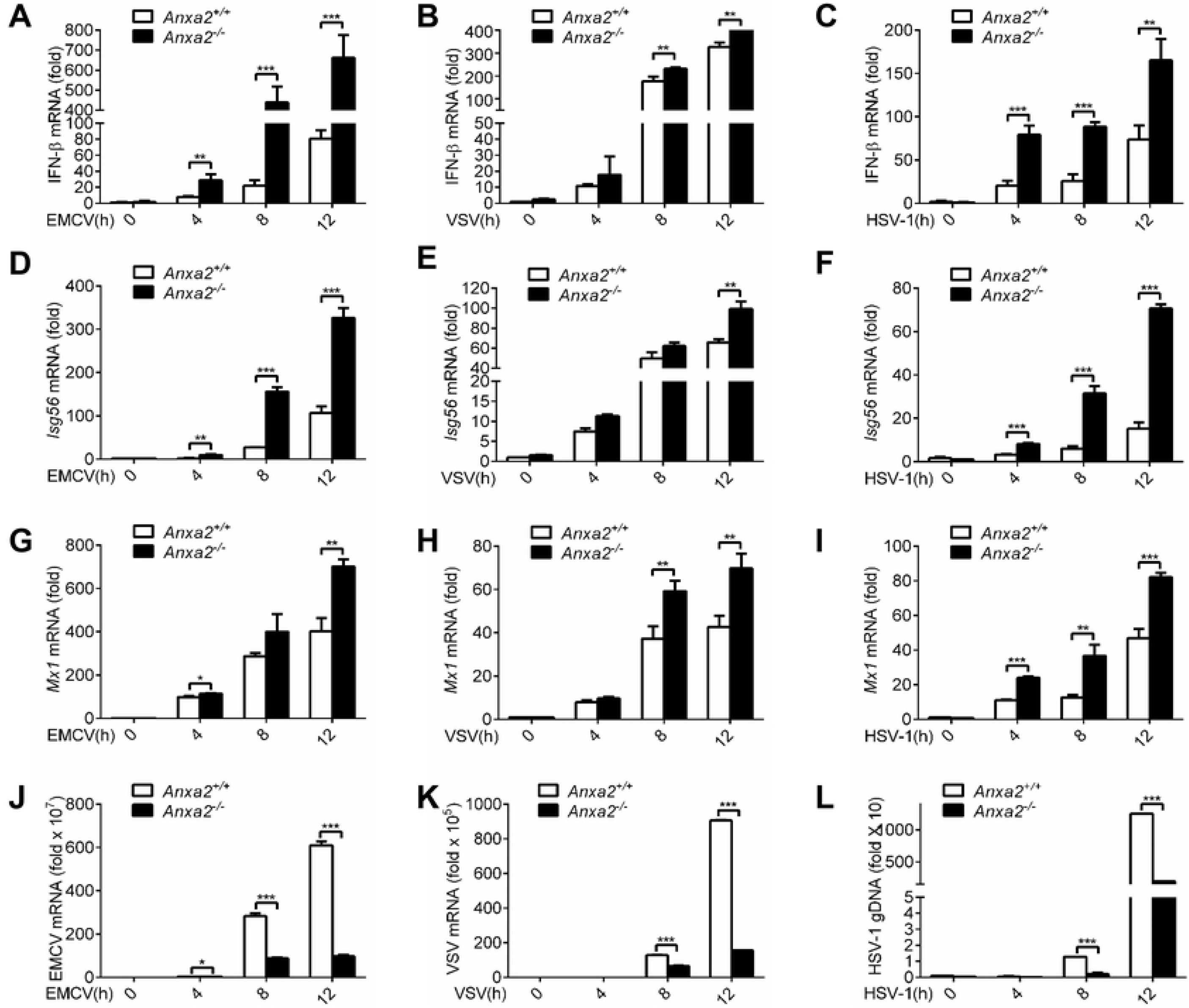

**Figure S5.**
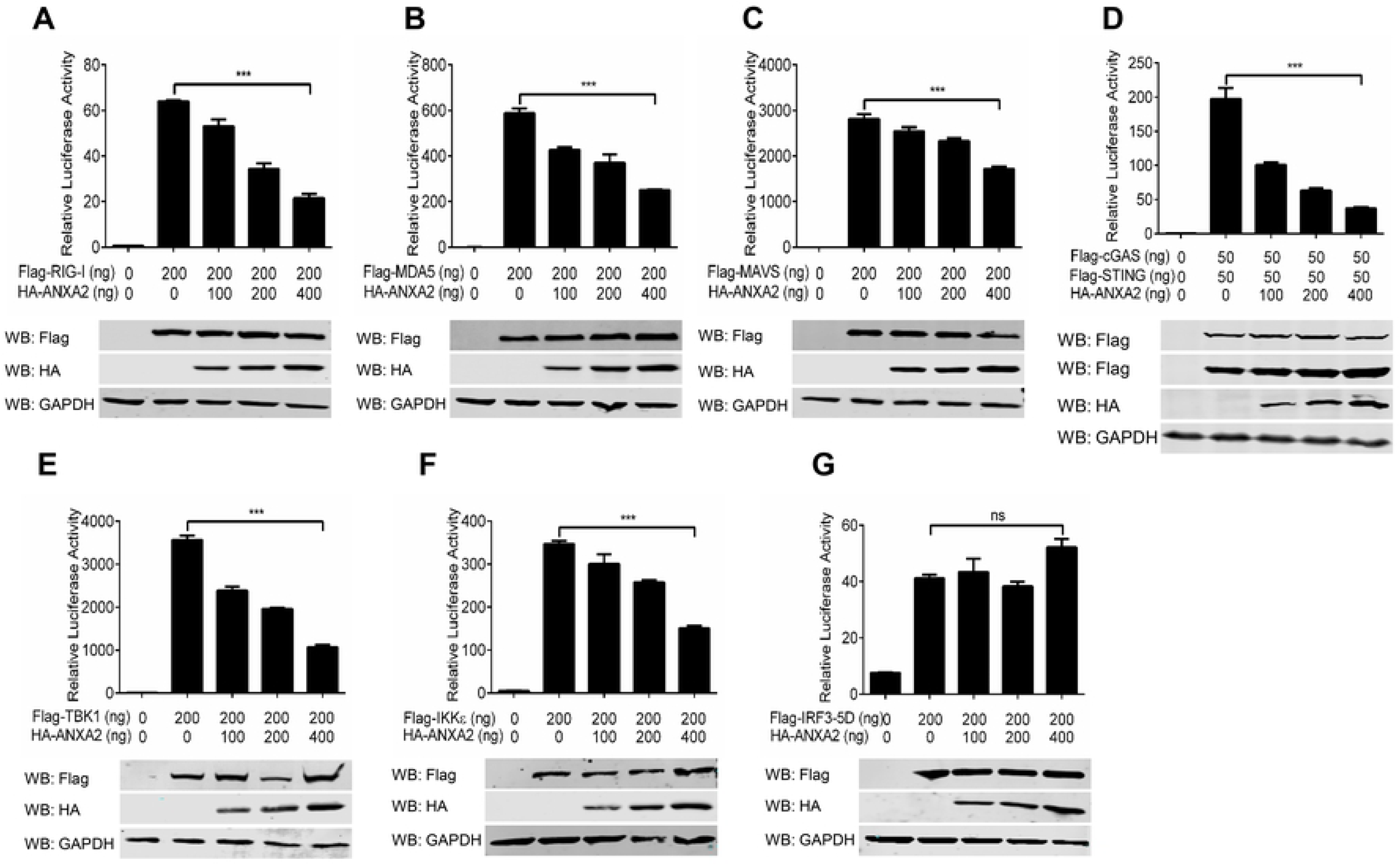

**Figure S6.**
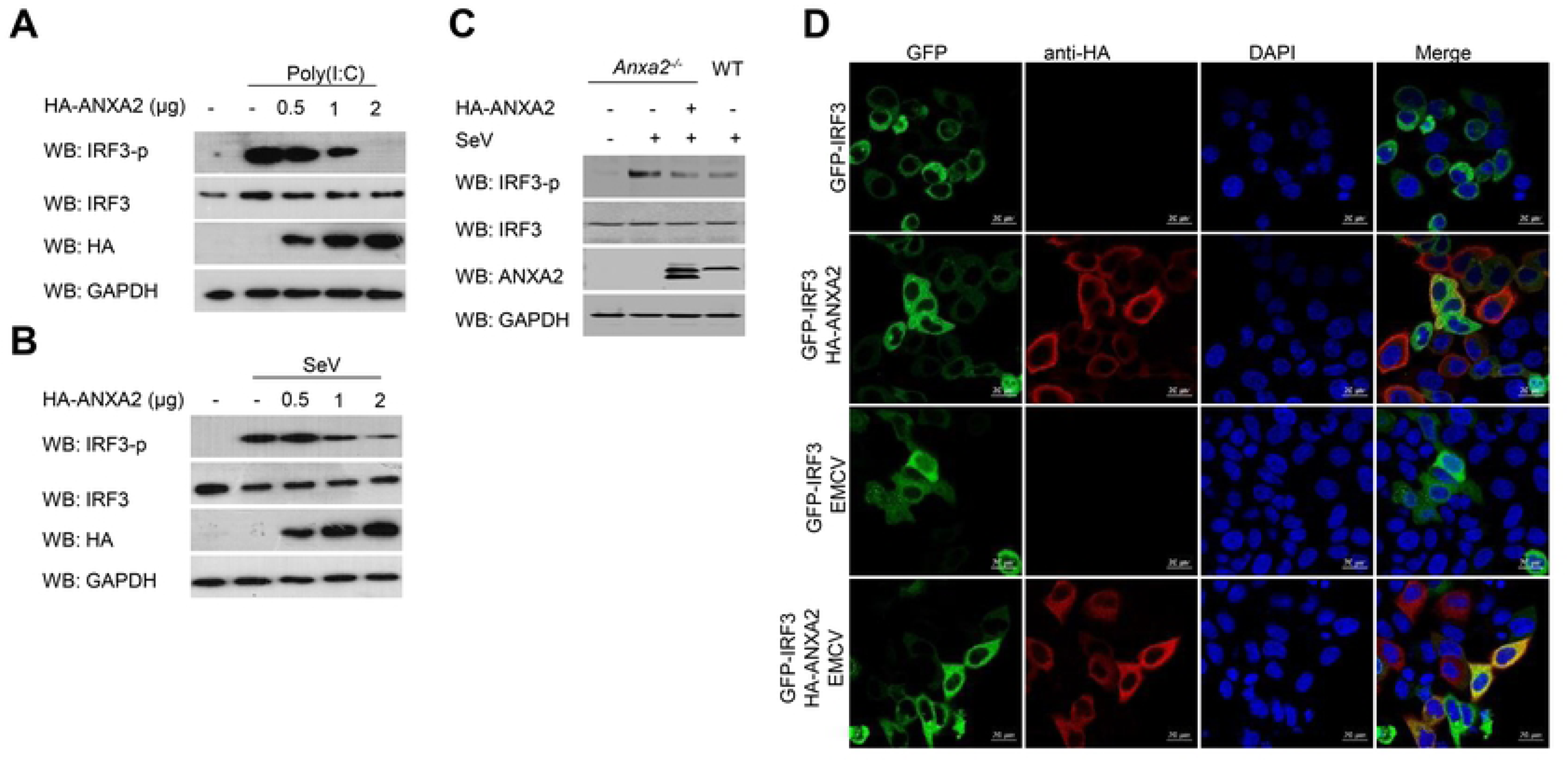

## Notes

### Competing Interest Statement

The authors have declared no competing interest.

## References

1. Kawai T, Akira S. The role of pattern-recognition receptors in innate immunity: update on Toll-like receptors. Nature immunology. 2010;11(5):373–84. Epub 2010/04/21. doi: 10.1038/ni.1863. PMID: 20404851.

2. Lupfer C, Kanneganti TD. Unsolved Mysteries in NLR Biology. Frontiers in immunology. 2013;4:285. Epub 2013/09/26. doi: 10.3389/fimmu.2013.00285. PMID: 24062750.

3. Yoneyama M, Kikuchi M, Natsukawa T, Shinobu N, Imaizumi T, Miyagishi M, et al. The RNA helicase RIG-I has an essential function in double-stranded RNA-induced innate antiviral responses. Nature immunology. 2004;5(7):730–7. Epub 2004/06/23. doi: 10.1038/ni1087. PMID: 15208624.

4. Scott I. The role of mitochondria in the mammalian antiviral defense system. Mitochondrion. 2010;10(4):316–20. Epub 2010/03/09. doi: 10.1016/j.mito.2010.02.005. PMID: 20206303; PubMed.

5. Seth RB, Sun L, Ea CK, Chen ZJ. Identification and characterization of MAVS, a mitochondrial antiviral signaling protein that activates NF-kappaB and IRF3. Cell. 2005;122(5):669–82. Epub 2005/08/30. doi: 10.1016/j.cell.2005.08.012. PMID: 16125763.

6. Takeuchi O, Akira S. Innate immunity to virus infection. Immunological reviews. 2009;227(1):75–86. Epub 2009/01/06. doi: 10.1111/j.1600-065X.2008.00737.x. PMID: 19120477.

7. Zhang X, Shi H, Wu J, Zhang X, Sun L, Chen C, et al. Cyclic GMP-AMP containing mixed phosphodiester linkages is an endogenous high-affinity ligand for STING. Molecular cell. 2013;51(2):226–35. Epub 2013/06/12. doi: 10.1016/j.molcel.2013.05.022. PMID: 23747010.

8. Yin Q, Tian Y, Kabaleeswaran V, Jiang X, Tu D, Eck MJ, et al. Cyclic di-GMP sensing via the innate immune signaling protein STING. Molecular cell. 2012;46(6):735–45. Epub 2012/06/19. doi: 10.1016/j.molcel.2012.05.029. PMID: 22705373.

9. Shu C, Yi G, Watts T, Kao CC, Li P. Structure of STING bound to cyclic di-GMP reveals the mechanism of cyclic dinucleotide recognition by the immune system. Nature structural & molecular biology. 2012;19(7):722–4. Epub 2012/06/26. doi: 10.1038/nsmb.2331. PMID: 22728658.

10. Li Y, Wilson HL, Kiss-Toth E. Regulating STING in health and disease. Journal of inflammation (London, England). 2017;14:11. Epub 2017/06/10. doi: 10.1186/s12950-017-0159-2. PMID: 28596706.

11. Ishikawa H, Barber GN. STING is an endoplasmic reticulum adaptor that facilitates innate immune signalling. Nature. 2008;455(7213):674–8. Epub 2008/08/30. doi: 10.1038/nature07317. PMID: 18724357.

12. Barber GN. STING: infection, inflammation and cancer. Nature reviews Immunology. 2015;15(12):760–70. Epub 2015/11/26. doi: 10.1038/nri3921. PMID: 26603901.

13. Liu S, Cai X, Wu J, Cong Q, Chen X, Li T, et al. Phosphorylation of innate immune adaptor proteins MAVS, STING, and TRIF induces IRF3 activation. Science (New York, NY). 2015;347(6227):aaa2630. Epub 2015/02/01. doi: 10.1126/science.aaa2630. PMID: 25636800.

14. Weaver BK, Kumar KP, Reich NC. Interferon regulatory factor 3 and CREB-binding protein/p300 are subunits of double-stranded RNA-activated transcription factor DRAF1. Molecular and cellular biology. 1998;18(3):1359–68. Epub 1998/03/06. doi: 10.1128/mcb.18.3.1359. PMID: 9488451.

15. Meylan E, Tschopp J, Karin M. Intracellular pattern recognition receptors in the host response. Nature. 2006;442(7098):39–44. Epub 2006/07/11. doi: 10.1038/nature04946. PMID: 16823444.

16. Yoneyama M, Suhara W, Fukuhara Y, Fukuda M, Nishida E, Fujita T. Direct triggering of the type I interferon system by virus infection: activation of a transcription factor complex containing IRF-3 and CBP/p300. The EMBO journal. 1998;17(4):1087–95. Epub 1998/03/28. doi: 10.1093/emboj/17.4.1087. PMID: 9463386.

17. Raynal P, Pollard HB. Annexins: the problem of assessing the biological role for a gene family of multifunctional calcium- and phospholipid-binding proteins. Biochimica et biophysica acta. 1994;1197(1):63–93. Epub 1994/04/05. doi: 10.1016/0304-4157(94)90019-1. PMID: 8155692.

18. Aliyu IA, Ling KH, Md Hashim N, Chee HY. Annexin A2 extracellular translocation and virus interaction: A potential target for antivirus-drug discovery. Reviews in medical virology. 2019;29(3):e2038. Epub 2019/02/13. doi: 10.1002/rmv.2038. PMID: 30746844.

19. Liu Y, Myrvang HK, Dekker LV. Annexin A2 complexes with S100 proteins: structure, function and pharmacological manipulation. British journal of pharmacology. 2015;172(7):1664–76. Epub 2014/10/11. doi: 10.1111/bph.12978. PMID: 25303710.

20. Réty S, Sopkova J, Renouard M, Osterloh D, Gerke V, Tabaries S, et al. The crystal structure of a complex of p11 with the annexin II N-terminal peptide. Nature structural biology. 1999;6(1):89–95. Epub 1999/01/14. doi: 10.1038/4965. PMID: 9886297.

21. Oh YS, Gao P, Lee KW, Ceglia I, Seo JS, Zhang X, et al. SMARCA3, a chromatin-remodeling factor, is required for p11-dependent antidepressant action. Cell. 2013;152(4):831–43. Epub 2013/02/19. doi: 10.1016/j.cell.2013.01.014. PMID: 23415230.

22. Koga R, Kubota M, Hashiguchi T, Yanagi Y, Ohno S. Annexin A2 Mediates the Localization of Measles Virus Matrix Protein at the Plasma Membrane. Journal of virology. 2018;92(10). Epub 2018/03/02. doi: 10.1128/jvi.00181-18. PMID: 29491166.

23. Chen L, Li X, Wang H, Hou P, He H. Annexin A2 gene interacting with viral matrix protein to promote bovine ephemeral fever virus release. Journal of veterinary science. 2020;21(2):e33. Epub 2020/04/02. doi: 10.4142/jvs.2020.21.e33. PMID: 32233139.

24. Sheng C, Liu X, Jiang Q, Xu B, Zhou C, Wang Y, et al. Annexin A2 is involved in the production of classical swine fever virus infectious particles. The Journal of general virology. 2015;96(Pt 5):1027–32. Epub 2015/01/17. doi: 10.1099/vir.0.000048. PMID: 25593157.

25. Saxena V, Lai CK, Chao TC, Jeng KS, Lai MM. Annexin A2 is involved in the formation of hepatitis C virus replication complex on the lipid raft. Journal of virology. 2012;86(8):4139–50. Epub 2012/02/04. doi: 10.1128/jvi.06327-11. PMID: 22301157.

26. Ma Y, Sun J, Gu L, Bao H, Zhao Y, Shi L, et al. Annexin A2 (ANXA2) interacts with nonstructural protein 1 and promotes the replication of highly pathogenic H5N1 avian influenza virus. BMC microbiology. 2017;17(1):191. Epub 2017/09/13. doi: 10.1186/s12866-017-1097-0. PMID: 28893180.

27. Chang XB, Yang YQ, Gao JC, Zhao K, Guo JC, Ye C, et al. Annexin A2 binds to vimentin and contributes to porcine reproductive and respiratory syndrome virus multiplication. Veterinary research. 2018;49(1):75. Epub 2018/07/29. doi: 10.1186/s13567-018-0571-5. PMID: 30053894.

28. Zerbe CM, Mouser DJ, Cole JL. Oligomerization of RIG-I and MDA5 2CARD domains. Protein science : a publication of the Protein Society. 2020;29(2):521–6. Epub 2019/11/08. doi: 10.1002/pro.3776. PMID: 31697400.

29. Zhu W, Li J, Zhang R, Cai Y, Wang C, Qi S, et al. TRAF3IP3 mediates the recruitment of TRAF3 to MAVS for antiviral innate immunity. The EMBO journal. 2019;38(18):e102075. Epub 2019/08/08. doi: 10.15252/embj.2019102075. PMID: 31390091.

30. Ishikawa H, Ma Z, Barber GN. STING regulates intracellular DNA-mediated, type I interferon-dependent innate immunity. Nature. 2009;461(7265):788–92. Epub 2009/09/25. doi: 10.1038/nature08476. PMID: 19776740.

31. Gerke V, Moss SE. Annexins: from structure to function. Physiological reviews. 2002;82(2):331–71. Epub 2002/03/28. doi: 10.1152/physrev.00030.2001. PMID: 11917092.

32. Waisman DM. Annexin II tetramer: structure and function. Molecular and cellular biochemistry. 1995;149-150:301–22. Epub 1995/08/01. doi: 10.1007/bf01076592. PMID: 8569746.

33. Pomerantz JL, Baltimore D. NF-kappaB activation by a signaling complex containing TRAF2, TANK and TBK1, a novel IKK-related kinase. The EMBO journal. 1999;18(23):6694–704. Epub 1999/12/03. doi: 10.1093/emboj/18.23.6694. PMID: 10581243.

34. Chariot A, Leonardi A, Muller J, Bonif M, Brown K, Siebenlist U. Association of the adaptor TANK with the I kappa B kinase (IKK) regulator NEMO connects IKK complexes with IKK epsilon and TBK1 kinases. The Journal of biological chemistry. 2002;277(40):37029–36. Epub 2002/07/23. doi: 10.1074/jbc.M205069200. PMID: 12133833.

35. Clark K, Peggie M, Plater L, Sorcek RJ, Young ER, Madwed JB, et al. Novel cross-talk within the IKK family controls innate immunity. The Biochemical journal. 2011;434(1):93–104. Epub 2010/12/09. doi: 10.1042/bj20101701. PMID: 21138416.

36. Clark K, Takeuchi O, Akira S, Cohen P. The TRAF-associated protein TANK facilitates cross-talk within the IkappaB kinase family during Toll-like receptor signaling. Proceedings of the National Academy of Sciences of the United States of America. 2011;108(41):17093–8. Epub 2011/09/29. doi: 10.1073/pnas.1114194108. PMID: 21949249.

37. Chau TL, Gioia R, Gatot JS, Patrascu F, Carpentier I, Chapelle JP, et al. Are the IKKs and IKK-related kinases TBK1 and IKK-epsilon similarly activated? Trends in biochemical sciences. 2008;33(4):171–80. Epub 2008/03/21. doi: 10.1016/j.tibs.2008.01.002. PMID: 18353649.

38. Rescher U, Zobiack N, Gerke V. Intact Ca(2+)-binding sites are required for targeting of annexin 1 to endosomal membranes in living HeLa cells. Journal of cell science. 2000;113 ( Pt 22):3931–8. Epub 2000/11/01. PMID: 11058080.

39. Alvarez-Martinez MT, Porte F, Liautard JP, Sri Widada J. Effects of profilin-annexin I association on some properties of both profilin and annexin I: modification of the inhibitory activity of profilin on actin polymerization and inhibition of the self-association of annexin I and its interactions with liposomes. Biochimica et biophysica acta. 1997;1339(2):331–40. Epub 1997/05/23. doi: 10.1016/s0167-4838(97)00018-6. PMID: 9187254.

40. Nilius B, Gerke V, Prenen J, Szücs G, Heinke S, Weber K, et al. Annexin II modulates volume-activated chloride currents in vascular endothelial cells. The Journal of biological chemistry. 1996;271(48):30631–6. Epub 1996/11/29. doi: 10.1074/jbc.271.48.30631. PMID: 8940038.

41. Díaz-Muñoz M, Hamilton SL, Kaetzel MA, Hazarika P, Dedman JR. Modulation of Ca2+ release channel activity from sarcoplasmic reticulum by annexin VI (67-kDa calcimedin). The Journal of biological chemistry. 1990;265(26):15894–9. Epub 1990/09/15. PMID: 2168425.

42. Naciff JM, Behbehani MM, Kaetzel MA, Dedman JR. Annexin VI modulates Ca2+ and K+ conductances of spinal cord and dorsal root ganglion neurons. The American journal of physiology. 1996;271(6 Pt 1):C2004–15. Epub 1996/12/01. doi: 10.1152/ajpcell.1996.271.6.C2004. PMID: 8997203.

43. Mussunoor S, Murray GI. The role of annexins in tumour development and progression. The Journal of pathology. 2008;216(2):131–40. Epub 2008/08/14. doi: 10.1002/path.2400. PMID: 18698663.

44. Madureira PA, Hill R, Miller VA, Giacomantonio C, Lee PW, Waisman DM. Annexin A2 is a novel cellular redox regulatory protein involved in tumorigenesis. Oncotarget. 2011;2(12):1075–93. Epub 2011/12/22. doi: 10.18632/oncotarget.375. PMID: 22185818.

45. Yee DS, Narula N, Ramzy I, Boker J, Ahlering TE, Skarecky DW, et al. Reduced annexin II protein expression in high-grade prostatic intraepithelial neoplasia and prostate cancer. Archives of pathology & laboratory medicine. 2007;131(6):902–8. Epub 2007/06/07. doi: 10.1043/1543-2165(2007)131[902:Raipei]2.0.Co;2. PMID: 17550317.

46. Scharf B, Clement CC, Wu XX, Morozova K, Zanolini D, Follenzi A, et al. Annexin A2 binds to endosomes following organelle destabilization by particulate wear debris. Nature communications. 2012;3:755. Epub 2012/03/29. doi: 10.1038/ncomms1754. PMID: 22453828.

47. Wang X, Shaw DK, Sakhon OS, Snyder GA, Sundberg EJ, Santambrogio L, et al. The Tick Protein Sialostatin L2 Binds to Annexin A2 and Inhibits NLRC4-Mediated Inflammasome Activation. Infection and immunity. 2016;84(6):1796–805. Epub 2016/04/06. doi: 10.1128/iai.01526-15. PMID: 27045038.

48. Bist P, Shu S, Lee H, Arora S, Nair S, Lim JY, et al. Annexin-A1 regulates TLR-mediated IFN-β production through an interaction with TANK-binding kinase 1. Journal of immunology (Baltimore, Md : 1950). 2013;191(8):4375–82. Epub 2013/09/21. doi: 10.4049/jimmunol.1301504. PMID: 24048896.

49. Yap GLR, Sachaphibulkij K, Foo SL, Cui J, Fairhurst AM, Lim LHK. Annexin-A1 promotes RIG-I-dependent signaling and apoptosis via regulation of the IRF3-IFNAR-STAT1-IFIT1 pathway in A549 lung epithelial cells. Cell death & disease. 2020;11(6):463. Epub 2020/06/17. doi: 10.1038/s41419-020-2625-7. PMID: 32541772.

50. Guo K, Lin X, Li Y, Qian W, Zou Z, Chen H, et al. Proteomic analysis of chicken embryo fibroblast cells infected with recombinant H5N1 avian influenza viruses with and without NS1 eIF4GI binding domain. Oncotarget. 2018;9(9):8350–67. Epub 2018/03/02. doi: 10.18632/oncotarget.23615. PMID: 29492200.

51. Takeuchi O, Akira S. MDA5/RIG-I and virus recognition. Current opinion in immunology. 2008;20(1):17–22. Epub 2008/02/15. doi: 10.1016/j.coi.2008.01.002. PMID: 18272355.

52. Xia P, Wang S, Gao P, Gao G, Fan Z. DNA sensor cGAS-mediated immune recognition. Protein & cell. 2016;7(11):777–91. Epub 2016/11/01. doi: 10.1007/s13238-016-0320-3. PMID: 27696330.

53. Uccellini MB, García-Sastre A. ISRE-Reporter Mouse Reveals High Basal and Induced Type I IFN Responses in Inflammatory Monocytes. Cell reports. 2018;25(10):2784–96.e3. Epub 2018/12/06. doi: 10.1016/j.celrep.2018.11.030. PMID: 30517866.

54. Zhang C, Shang G, Gui X, Zhang X, Bai XC, Chen ZJ. Structural basis of STING binding with and phosphorylation by TBK1. Nature. 2019;567(7748):394–8. Epub 2019/03/08. doi: 10.1038/s41586-019-1000-2. PMID: 30842653.

55. Li P, Zhu Z, Cao W, Yang F, Ma X, Tian H, et al. Dysregulation of the RIG-I-like Receptor Pathway Signaling by Peste des Petits Ruminants Virus Phosphoprotein. Journal of immunology (Baltimore, Md : 1950). 2021;206(3):566–79. Epub 2021/01/01. doi: 10.4049/jimmunol.2000432. PMID: 33380495.

56. Kumar KP, McBride KM, Weaver BK, Dingwall C, Reich NC. Regulated nuclear-cytoplasmic localization of interferon regulatory factor 3, a subunit of double-stranded RNA-activated factor 1. Molecular and cellular biology. 2000;20(11):4159–68. Epub 2000/05/11. doi: 10.1128/mcb.20.11.4159-4168.2000. PMID: 10805757.

57. Wang P, Zhao W, Zhao K, Zhang L, Gao C. TRIM26 negatively regulates interferon-β production and antiviral response through polyubiquitination and degradation of nuclear IRF3. PLoS pathogens. 2015;11(3):e1004726. Epub 2015/03/13. doi: 10.1371/journal.ppat.1004726. PMID: 25763818.

58. Zhang K, Zhang Y, Xue J, Meng Q, Liu H, Bi C, et al. DDX19 Inhibits Type I Interferon Production by Disrupting TBK1-IKKε-IRF3 Interactions and Promoting TBK1 and IKKε Degradation. Cell reports. 2019;26(5):1258–72.e4. Epub 2019/01/31. doi: 10.1016/j.celrep.2019.01.029. PMID: 30699353.

59. Gerke V, Creutz CE, Moss SE. Annexins: linking Ca2+ signalling to membrane dynamics. Nature reviews Molecular cell biology. 2005;6(6):449–61. Epub 2005/06/02. doi: 10.1038/nrm1661. PMID: 15928709.

60. Dallacasagrande V, Hajjar KA. Annexin A2 in Inflammation and Host Defense. Cells. 2020;9(6). Epub 2020/06/25. doi: 10.3390/cells9061499. PMID: 32575495.

61. Zhang S, Yu M, Guo Q, Li R, Li G, Tan S, et al. Annexin A2 binds to endosomes and negatively regulates TLR4-triggered inflammatory responses via the TRAM-TRIF pathway. Scientific reports. 2015;5:15859. Epub 2015/11/04. doi: 10.1038/srep15859. PMID: 26527544.

62. Stukes S, Coelho C, Rivera J, Jedlicka AE, Hajjar KA, Casadevall A. The Membrane Phospholipid Binding Protein Annexin A2 Promotes Phagocytosis and Nonlytic Exocytosis of Cryptococcus neoformans and Impacts Survival in Fungal Infection. Journal of immunology (Baltimore, Md : 1950). 2016;197(4):1252–61. Epub 2016/07/03. doi: 10.4049/jimmunol.1501855. PMID: 27371724.

63. Ran FA, Hsu PD, Wright J, Agarwala V, Scott DA, Zhang F. Genome engineering using the CRISPR-Cas9 system. Nature protocols. 2013;8(11):2281–308. Epub 2013/10/26. doi: 10.1038/nprot.2013.143. PMID: 24157548.

